# Compatibility logic of human enhancer and promoter sequences

**DOI:** 10.1101/2021.10.23.462170

**Authors:** Drew T. Bergman, Thouis R. Jones, Vincent Liu, Layla Siraj, Helen Y. Kang, Joseph Nasser, Michael Kane, Tung H. Nguyen, Sharon R. Grossman, Charles P. Fulco, Eric S. Lander, Jesse M. Engreitz

## Abstract

Gene regulation in the human genome is controlled by distal enhancers that activate specific nearby promoters. One model for the specificity of enhancer-promoter regulation is that different promoters might have sequence-encoded preferences for distinct classes of enhancers, for example mediated by interacting sets of transcription factors or cofactors. This “biochemical compatibility” model has been supported by observations at individual human promoters and by genome-wide measurements in *Drosophila*. However, the degree to which human enhancers and promoters are intrinsically compatible or specific has not been systematically measured, and how their activities combine to control RNA expression remains unclear. To address these questions, we designed a high-throughput reporter assay called enhancer x promoter (ExP) STARR-seq and applied it to examine the combinatorial compatibilities of 1,000 enhancer and 1,000 promoter sequences in human K562 cells. We identify a simple logic for enhancer-promoter compatibility – virtually all enhancers activated all promoters by similar amounts, and intrinsic enhancer and promoter activities combine multiplicatively to determine RNA output (*R^2^*=0.82). In addition, two classes of enhancers and promoters showed subtle preferential effects. Promoters of housekeeping genes contained built-in activating sequences, corresponding to motifs for factors such as GABPA and YY1, that correlated with both stronger autonomous promoter activity and enhancer activity, and weaker responsiveness to distal enhancers. Promoters of context-specific genes lacked these motifs and showed stronger responsiveness to enhancers. Together, this systematic assessment of enhancer-promoter compatibility suggests a multiplicative model tuned by enhancer and promoter class to control gene transcription in the human genome.

## Introduction

The extent to which distal enhancers might activate specific types of promoters has been an outstanding question in human gene regulation. Since their initial discovery, enhancers have been defined in part based on their ability to activate multiple non-cognate promoter sequences^1, 2^. High-throughput reporter assays have now confirmed that many enhancer sequences derived from the human genome have the capability to activate various human, viral, and synthetic promoters^3–9^.

Yet, other observations have suggested that enhancers and promoters have some degree of intrinsic specificity. Early studies identified individual examples where particular enhancers or cofactors showed stronger activation with certain core promoters^10–15^. More recently, in *Drosophila*, studies using high-throughput reporter assays revealed that developmental and housekeeping gene promoters show >10-fold preferences for different classes of genomic enhancers^16^, have differing levels of sequence-encoded responsiveness to enhancer activation^17^, and respond differently to recruitment of various transcriptional cofactors^18^. Together, these studies have suggested a ‘biochemical compatibility’ model where different enhancers might have an intrinsic preference for activating different promoter sequences based on the transcription factors and cofactors they can recruit^19, 20^.

Despite these advances, the biochemical compatibility model has not been systematically tested for human enhancers and promoters. As such, it remains unclear whether compatibility classes of enhancers and promoters exist in the human genome, and, if so, how their activities combine and how such specificity is encoded.

### High-throughput measurements of enhancer-promoter compatibility

To investigate these questions, we developed an assay called enhancer x promoter (ExP) STARR-seq to test the ability of ∼1,000 candidate enhancers to activate ∼1,000 promoters. In this assay, we synthesize pools of enhancer and promoter sequences (here, 264-bp) and clone them in all pairwise combinations located ∼340-bp apart in the revised human STARR-seq plasmid-based reporter vector (Fig. 1a**/**S1a)^8^. In STARR-seq assays, the enhancer sequence is transcribed and quantified using targeted RNA-seq to determine the level of expression of each plasmid^4^. For ExP STARR-seq, we introduce a unique 16-bp “plasmid barcode” adjacent to the enhancer sequence that allows us to determine which reporter transcripts are produced from which enhancer-promoter pairs. We transiently transfect this pool of plasmids into cells, measure the level of reporter transcripts produced, and calculate “STARR-seq expression” as the amount of RNA normalized to DNA input for each plasmid. This approach allows us to quantitatively measure the expression of hundreds of thousands of combinations of enhancer and promoter sequences, estimate the activities of individual enhancers and promoters, and test their compatibilities (see Methods).

**Fig. 1.**
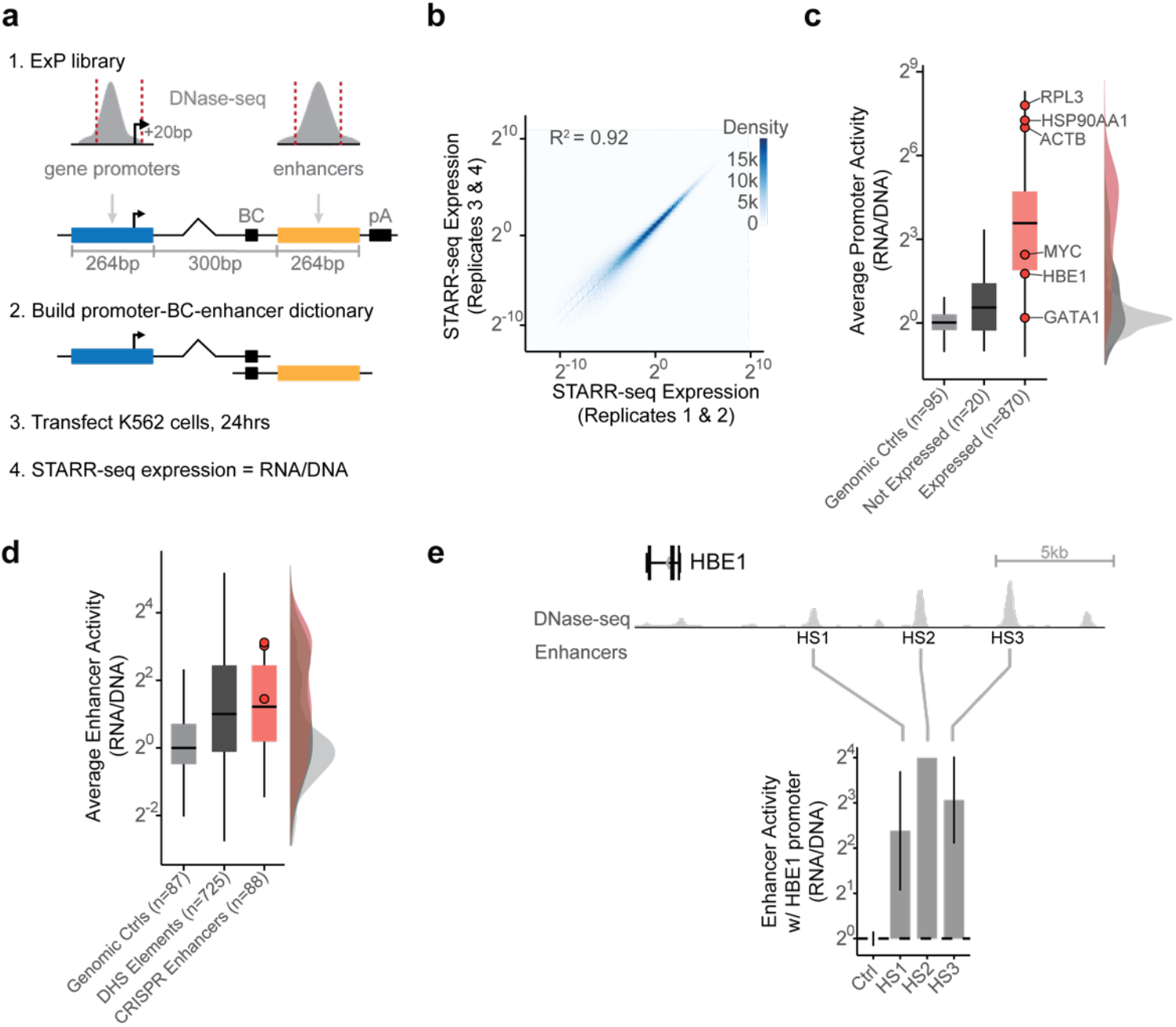
Enhancer x Promoter STARR-seq. **a.** ExP STARR-seq method for measuring the activities of enhancer and promoter sequences and testing their compatibilities. 264-bp sequences are selected and cloned in all pairwise combinations into the promoter and enhancer positions of a plasmid vector, together with a plasmid barcode (BC). We build a dictionary linking promoter-BC-enhancer triplets via sequencing (see Fig. S1a). We then transfect the ExP STARR-seq plasmid pool into cells, where the promoter sequence on a given plasmid initiates transcription of a polyadenylated RNA containing the plasmid barcode and enhancer. We sequence these RNAs and calculate STARR-seq expression as the frequency of RNAs observed for each plasmid normalized by the frequency of that plasmid in the input DNA plasmid pool. **b.** Correlation of ExP STARR-seq expression between biological replicate experiments, calculated for individual enhancer-promoter pairs with unique plasmid barcodes. Axes represent the average STARR-seq expression (RNA/DNA) of two biological replicates. Density: number of enhancer-promoter plasmids. **c.** Average promoter activity (STARR-seq expression when paired with random genomic controls in the enhancer position) of promoter sequences derived from random genomic controls (set at 0), genes not expressed in K562s, and all other gene promoters. Box is median and interquartile range, whiskers are +/- 1.5 x IQR. **d.** Average enhancer activity (STARR-seq expression of plasmids containing a given enhancer averaged across all promoters) of enhancer sequences derived from random genomic controls, accessible elements, and genomic enhancers validated in CRISPR experiments. Box and whiskers as in (**c**). Red dots represent three enhancers near *HBE1* (see panel **e**). **e.** Sequences derived from three genomic enhancers that regulate *HBE1* in the genome (HS1-HS3) activate the *HBE1* promoter in ExP STARR-seq. Ctrl: Average of 44 random genomic control sequences in the enhancer position that passed thresholds (see Methods). Error bars: 95% CI across plasmid barcodes, n=110 (ctrl), 2 (HS1), 1 (HS2), 5 (HS3).

Hereafter, for clarity, we use the terms “enhancer sequences” and “promoter sequences” to refer to sequences cloned into the enhancer and promoter positions in the ExP STARR-seq assay, and “genomic enhancers” and “genomic promoters” to refer to the corresponding elements in the genome.

We applied ExP STARR-seq to examine the combinatorial activities of 1,000 enhancer and 1,000 promoter sequences (**Table S1**, **Table S2**) in K562 erythroleukemia cells, which have been deeply profiled by the ENCODE Project^21^ and where we have previously collected data about which genomic enhancers regulate which genomic promoters using CRISPR interference (CRISPRi) screens^22^. Here, we selected promoter sequences to include (i) 65 genes studied in prior CRISPR screens; (ii) 735 additional genes sampled from across the genome to span a range of transcriptional activity (based on precision run-on sequencing (PRO-seq) data in K562 cells); and (iii) 200 control sequences including random genomic control sequences that are not accessible by ATAC-seq, and dinucleotide shuffled sequences (Fig. S1a, see Methods). The promoter sequences were chosen to include approximately 20-bp downstream of the genomic transcription start site (as observed in capped analysis of gene expression (CAGE) data), and ∼242-bp upstream (264 bp total, see Methods). In the enhancer position of ExP STARR-seq, we included (i) 131 accessible genomic elements we previously tested by CRISPRi; (ii) 669 other accessible genomic elements selected to span a range of quantitative H3K27ac and DNase-seq signals (centered on the summit of the DNase-seq peak); and (iii) 200 controls including random genomic control sequences and dinucleotide shuffled sequences (Fig. S1a, See Methods).

We cloned these 1,000 enhancer and 1,000 promoter sequences in all pairwise combinations, transfected the plasmid pool into K562 cells in 4 biological replicates of 50 million cells each, and sequenced each STARR-seq RNA and input DNA library to a depth of at least 2.6 billion and 470 million reads, respectively. We focused our analysis on the 604,268 enhancer-promoter pairs where we obtained good coverage (see Methods). STARR-seq expression (RNA/DNA) varied over six orders of magnitude, and was highly reproducible, when comparing expression for individual plasmid barcodes between biological replicates (*R*^2^ = 0.92, Fig. 1b), when comparing expression for an enhancer-promoter pair averaged across plasmid barcodes between biological replicates (*R*^2^ = 0.92), and when comparing expression for different plasmid barcodes for a given enhancer-promoter pair (*R*^2^ = 0.62, Fig. S1c-d, see Methods).

As expected, promoter sequences showed a very large (>1,500-fold) dynamic range of STARR-seq expression when paired with random genomic sequences in the enhancer position (“average promoter activity”). The strongest promoters in the dataset corresponded to housekeeping genes such as *RPL3*, *HSP90AA1*, and *ACTB*, and the weakest promoters included shuffled control sequences and non-expressed genes in K562 cells (Fig. 1c). Enhancer sequences also showed a wide (682-fold) range of STARR-seq expression in the dataset when averaged across promoters (“average enhancer activity”), and were on average 2-fold more active than random genomic control sequences (Fig. 1d). Enhancer and promoter activity from ExP STARR-seq were correlated with biochemical features of activity at the corresponding genomic elements, including with levels of chromatin accessibility, H3K27 acetylation, and nascent gene and eRNA transcription (Fig. S1e).

We also found that sequences derived from known genomic enhancers activated their cognate promoters in the ExP STARR-seq assay. For example, we included 3 enhancers in the beta-like globin locus control region (HS1-HS3) that are known to coordinate expression of hemoglobin subunits during erythrocyte development^23, 24^ and where CRISPRi perturbations in K562 cells reduce the expression of hemoglobin subunit epsilon 1 (*HBE1*) by 10-86%^25, 26^. In ExP STARR-seq, each of these enhancers activated the *HBE1* promoter (by 5.21-15.9-fold versus random genomic controls, Fig. 1e). Similarly, an enhancer that we previously showed to regulate *GATA1* and *HDAC6* in the genome^27^ led to 6.76 and 6.87-fold activation of the *GATA1* and *HDAC6* promoters in ExP STARR-seq, respectively (Fig. S1f).

Taken together, these results show that ExP-STARR-seq produces quantitative and reproducible measurements of enhancer and promoter sequence activity over a large dynamic range.

### Enhancer and promoter sequences are broadly compatible

We used this ExP STARR-seq dataset to test whether specific enhancers activate specific promoters. Surprisingly, virtually all active enhancer sequences activated all promoter sequences by similar amounts. For example, for a small subset of 5 enhancers and 5 promoters, each with good coverage in the assay (median = 27 plasmid barcodes per pair), while the promoters spanned a 5.62-fold range of activities, the enhancers activated each promoter similarly (Fig. 2a-b). More generally, enhancers activated most promoters by similar amounts, with an average Spearman correlation across all pairs of promoters = 0.81 (Fig. 2c,e, S2a), and pairs of enhancers showed similar proportional activation of promoters, with an average Spearman = 0.72 (Fig. 2d,f, S2b). These observations indicate that, in this STARR-seq assay, there is broad compatibility between individual enhancer and promoter sequences — a striking difference from previous observations in *Drosophila*^16, 17^.

**Fig. 2.**
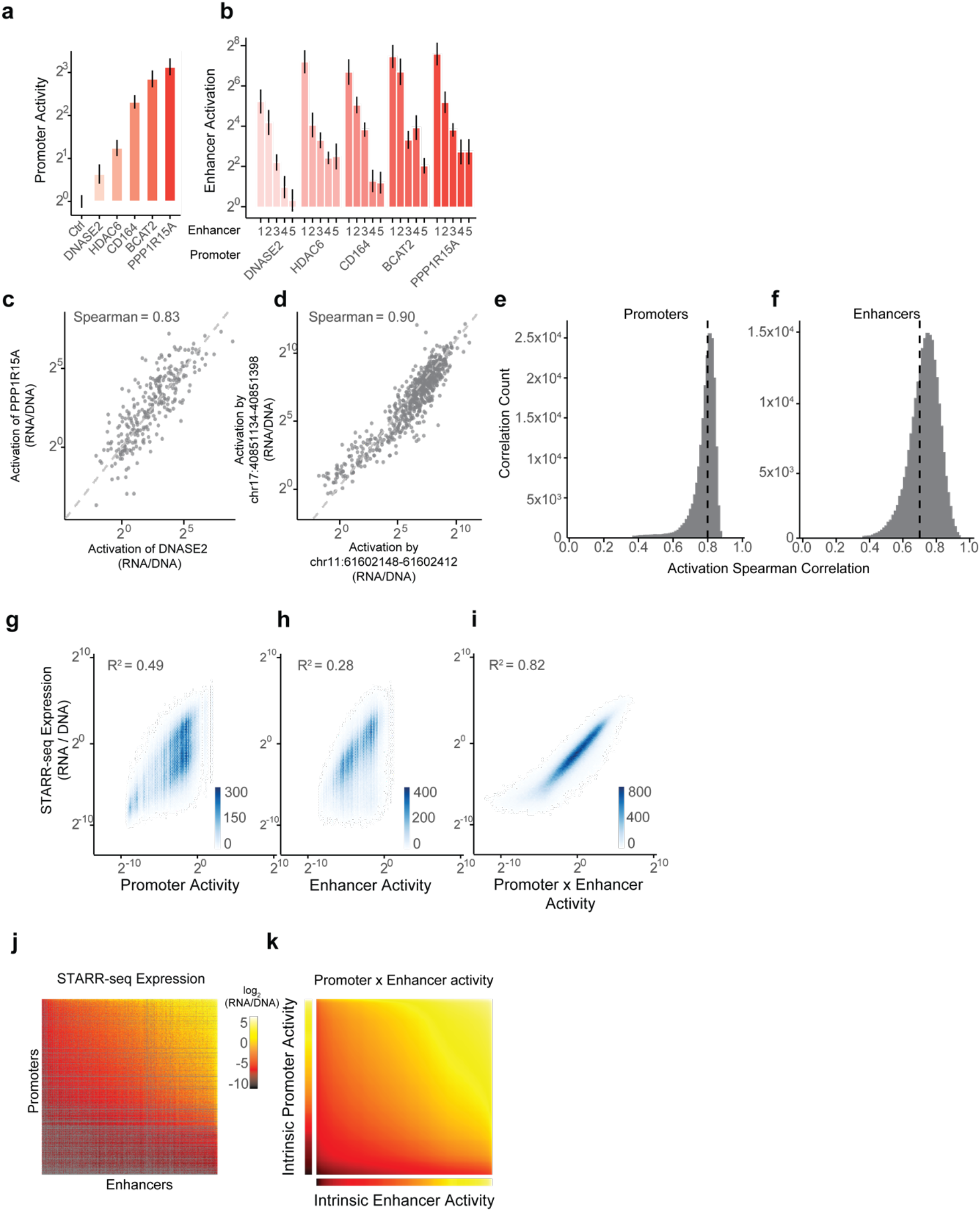
Enhancer and promoter activities combine multiplicatively. **a.** Intrinsic promoter activity (expression versus random genomic controls in enhancer position) of five selected promoters. Error bars: 95% CI across plasmid barcodes (n=54-79). **b.** Activation (expression versus random genomic controls in enhancer position) of 5 selected promoters by 5 selected enhancers (1 = chr11:61602148-61602412, 2 = chr19:49467061-49467325, 3 = chrX:48641342-48641606, 4 = chr19:12893216-12893480, 5 = chr17:40851134-40851398). Error bars: 95% CI across plasmid barcodes (n=12-56). **c.** Correlation of enhancer activation for PPP1R15A and DNASE2 promoters. Each point is a shared enhancer sequence. **d.** Correlation of enhancer activation by chr17:40851134-40851398 and chr11:61602148-61602412 enhancers. Each point is a shared promoter sequence. **e.** Distribution of pairwise correlations of enhancer activation between promoter sequences, as in (**c**). black dotted line = mean Spearman correlation. **f.** Distribution of pairwise correlations of promoter activation between enhancer sequences, as in (**d**). Black dotted line = mean Spearman correlation. **g-i.** Correlation of ExP STARR-seq expression with intrinsic promoter activity (**g**), intrinsic enhancer activity (**h**), and the product of intrinsic promoter and enhancer activities (**i**). Density color scale: number enhancer-promoter pairs. **j.** Heatmap of ExP STARR-seq expression across all pairs of promoter (vertical) and enhancer sequences (horizontal). Axes are sorted by intrinsic promoter and enhancer activities. Grey: missing data. **k.** Heatmap representing the multiplication of intrinsic promoter activity (vertical) with intrinsic enhancer activity (horizontal) from the Poisson model.

### Enhancer and promoter activities combine approximately multiplicatively

This pattern of effects — where enhancers showed similar fold-activation across many promoters, and promoters showed similar levels of activation by many enhancers — suggested that intrinsic enhancer and promoter activities combine multiplicatively to produce the RNA output in STARR-seq. To quantify this, we correlated expression in the STARR-seq assay with intrinsic enhancer activity, intrinsic promoter activity, and the multiplicative product of intrinsic enhancer and promoter activities.

To do so, we fit the following Poisson count model:

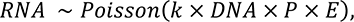

where *RNA* is RNA reads counts per plasmid, *DNA* is DNA read counts per plasmid, *P* is the intrinsic promoter activity, *E* is intrinsic enhancer activity, and *k* is a free intercept term used to scale the activities of promoters, enhancers, and their pairings relative to the average of random genomic control sequences (see Methods). This multiplicative model assumes that there is no sequence or biochemical specificity between individual pairs of enhancers and promoters, and that differences in expression are solely due to differences in intrinsic enhancer and promoter activities. Hereafter, we define “intrinsic enhancer activity” and “intrinsic promoter activity” as the fits from this model, which yield similar estimates to the “average activities” calculated above (Fig. S2c,d) but better account for missing data and counting noise (see Methods). These estimates of activity were reproducible across replicate experiments and when comparing non-overlapping plasmid barcodes (Fig. S2e,f).

Intrinsic promoter activity alone explained 49% of the variance in STARR-seq expression across all enhancer-promoter pairs (correlation with log_2_ STARR-seq expression in pairs with at least 2 plasmid barcodes, Fig. 2g), and intrinsic enhancer activity alone explained 28% of the variance (Fig. 2h). The multiplicative combination of intrinsic promoter and enhancer activities explained 82% of the total variance (Fig. 2i-k).

To confirm that this multiplicative relationship was not due to the specific design of our ExP STARR-seq assay, we cloned 7 enhancers from the *MYC* locus (1.0-2.2 kb) and 5 promoter sequences (138-908 bp, including the promoters of *MYC* and other nearby genes) in all combinations into a different reporter plasmid in which the enhancer is located 1 kb upstream of the promoter, and measured the expression of these constructs using a luciferase reporter assay (Fig. S2g, **Table S3**). Again, despite a range of intrinsic promoter activities (Fig. S2h), all enhancer sequences activated all promoter sequences by a similar fold-change, and a multiplicative function of enhancer and promoter activities explained 78% of the total variance in the measurements (Fig. S2i).

Thus, RNA expression in these reporter assays represents, to a first approximation, the multiplicative product of intrinsic enhancer activity and intrinsic promoter activity.

### Two functional classes of enhancer and promoter sequences

Although we did not observe a strong degree of specificity among enhancer and promoter sequences, we asked whether there might exist classes with more subtle, quantitative preferences. To do so, we calculated, for each enhancer-promoter pair, its deviation from the multiplicative enhancer x promoter model (observed STARR-seq expression versus the product of intrinsic enhancer activity and intrinsic promoter activity, see Methods). We identified two clusters of enhancer sequences (E1 and E2, n=126 and 290 respectively) that showed differential effects with respect to two sets of promoter sequences (P1 and P2, n=192 and 391 respectively) (Fig. 3a). In particular, E1 enhancer sequences activated P1 promoters more strongly than P2 promoters (by 1.93-fold, *P* = 4.19e-08, *t*-test), whereas E2 enhancer sequences activated promoters in both clusters approximately equally (1.05-fold stronger for P2 versus P1, *P* = 0.424, *t*-test; Fig. 3b). These sets of enhancers and promoters appeared to represent extremes of a graded scale: promoter responsiveness to E1 vs E2 enhancer sequences varied over a ∼3-fold range (Fig. 3c, Fig. S3d, Fig. S4b), and enhancer activation of P1 vs P2 promoters varied over a ∼2-fold range (Fig. 3d, Fig. S3e, Fig. S4a). Cluster assignments were highly stable to down-sampling of promoter and enhancer sequences (Fig. S3g, see Methods). Additional clusters (P0 and E0) contained sequences with very weak activity and/or missing data, and were excluded from further analysis (Fig. S3a-c).

**Fig. 3.**
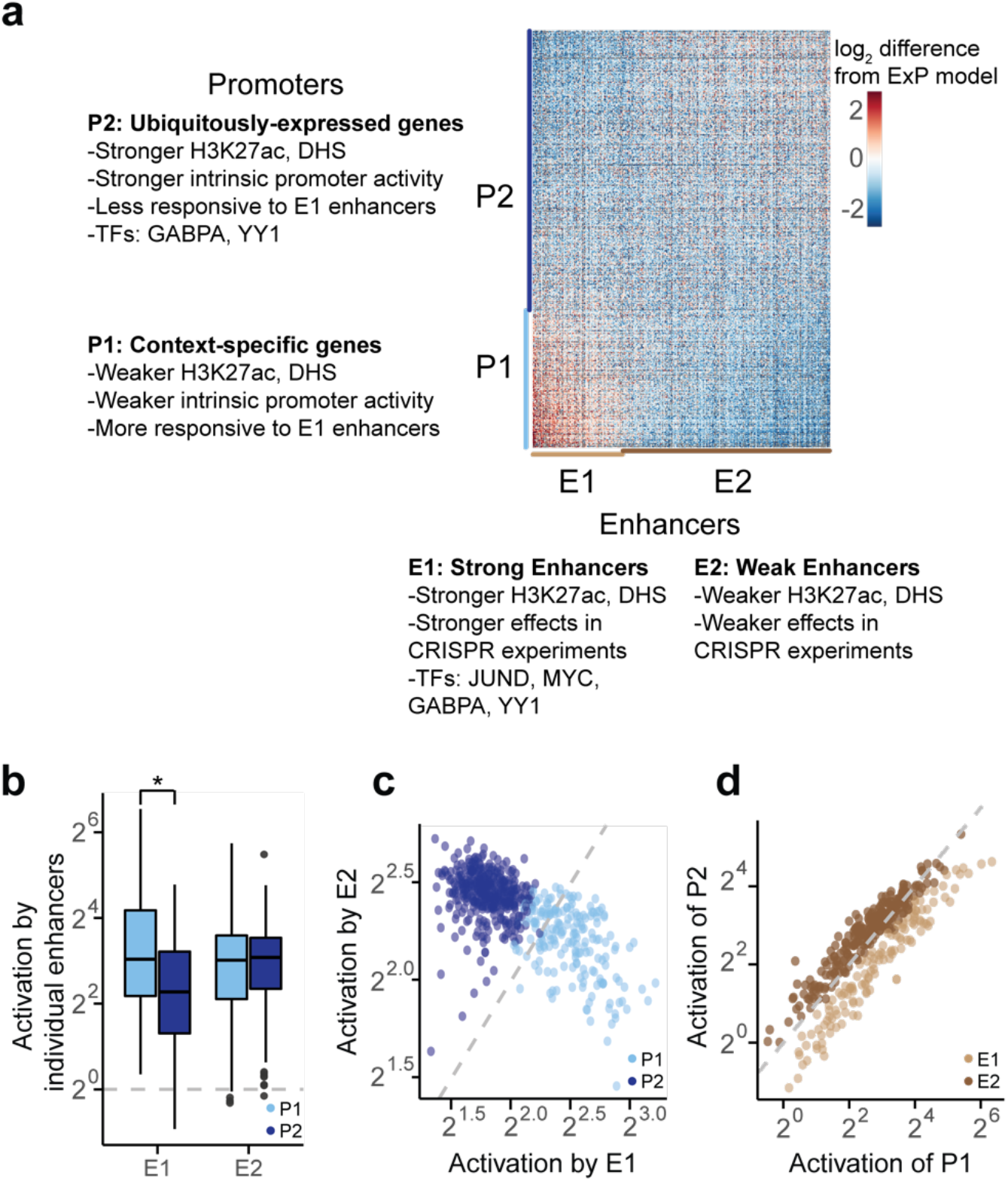
Compatibility classes of enhancers and promoters. **a.** Heatmap of deviations in enhancer-promoter STARR-seq expression from a multiplicative enhancer-promoter model (color scale: fold-difference between observed expression versus expression predicted by multiplicative model; gray: missing data). Vertical axis: promoter sequences grouped by class and sorted by responsiveness to E1 vs. E2 (see **b**); horizontal axis: enhancer sequences grouped by class and sorted by activation of P1 vs. P2 (see **c**). **b.** Activation of P1 vs P2 promoters by E1 and E2 enhancer sequences (equivalently: Responsiveness to E1 vs E2 enhancer sequences). Boxes are median and interquartile range, whiskers are +/- 1.5*IQR. \**P*-value = 4.2 x 10^-8^, two-sample *t*-test. **c.** For each promoter, the average activation by (responsiveness to) E1 enhancer sequences (x-axis) versus the average activation by E2 enhancer sequences (y-axis). P1 promoters (light blue) are activated more strongly by E1 versus E2 enhancers. **d.** For each enhancer, the average fold-activation when paired with P1 promoters (*x-*axis) versus P2 promoters (*y*-axis). E1 enhancers (light brown) more strongly activate P1 promoters.

Together, these observations identify 2 classes of enhancer sequences and 2 classes of promoter sequences with subtle quantitative differences in compatibility. Accordingly, we next sought to characterize these classes of enhancer and promoter sequences and understand how such preferential effects might be encoded.

### Classes of enhancer sequences correspond to strong and weak genomic enhancers

To characterize the two classes of ExP STARR-seq enhancer sequences, we compared the classes with respect to biochemical features of their corresponding elements in the genome, sequence motifs, effects in CRISPR experiments, and other features.

E1 and E2 classes showed biochemical features of strong and weak genomic enhancers, respectively. The features most strongly associated with E1 versus E2 sequences in the genome included H3K27ac, DNase I hypersensitivity, AP-1 factor binding (JUN, ATF3), and other known activating transcription factors (Fig. 3a, Fig. S5a-b, **Table S4**). E2 sequences in the genome were also DNase accessible and sometimes bound these factors, but to a significantly lesser degree (Fig. S7). Consistent with these observations, E1 sequences had stronger effects on gene expression in CRISPR perturbation experiments, even when controlling for 3D contact with the target gene (Fig. S5c). While E1 sequences were more likely to be predicted to be enhancers in K562 cells (94% of E1 predicted to regulate a gene by the Activity-by-Contact (ABC) model, versus 49% of E2), both classes contained a large fraction of sequences predicted to be an enhancer in another cell type (90% of E1 and 70% of E2), suggesting that some E2 genomic elements may act as strong enhancers in other cell types.

These observations suggest that the differences in how these classes of enhancer sequences activate different promoters in ExP-STARR-seq could be related to their ability to recruit activating transcription factors (see below). We note that, despite these clear differences in genomic activity, the two classes of enhancer sequences showed, on average, similar levels of activity in the ExP-STARR-seq assay (Fig. S3b). This may reflect previous observations that the episomal STARR-seq assay often detects activity for sequences that do not appear to be active in their endogenous chromosomal context^8, 28^.

### Classes of promoter sequences correspond to constitutive versus enhancer-responsive genes

The two classes of promoter sequences also showed striking differences in their functional annotations, intrinsic promoter activity, and responsiveness to enhancers in the genome.

We found that P2 promoter sequences were primarily derived from ubiquitously expressed genes (often called “housekeeping” genes), whereas P1 promoters corresponded to cell-type- or context-specific genes. For example, P2 promoters included beta actin (*ACTB*), all 37 tested ribosomal subunits (*e.g.*, *RPL13*, *RPS11*), components of the electron transport chain (*e.g.*, *NDUFA2*, *ATP5B*), and others (**Table S1**). In contrast, P1 promoters included erythroid-specific genes (*e.g.*, 3 hemoglobin genes, ferritin light chain (*FTL*)), context-specific transcription factors (*e.g.*, *KLF1*, *JUNB*, *REL*), and genes that are expressed in many cell types but at different levels, such as *MYC*. P1 and P2 promoters were associated with developmental and housekeeping gene ontology terms, respectively (Fig. 4a).

**Fig. 4.**
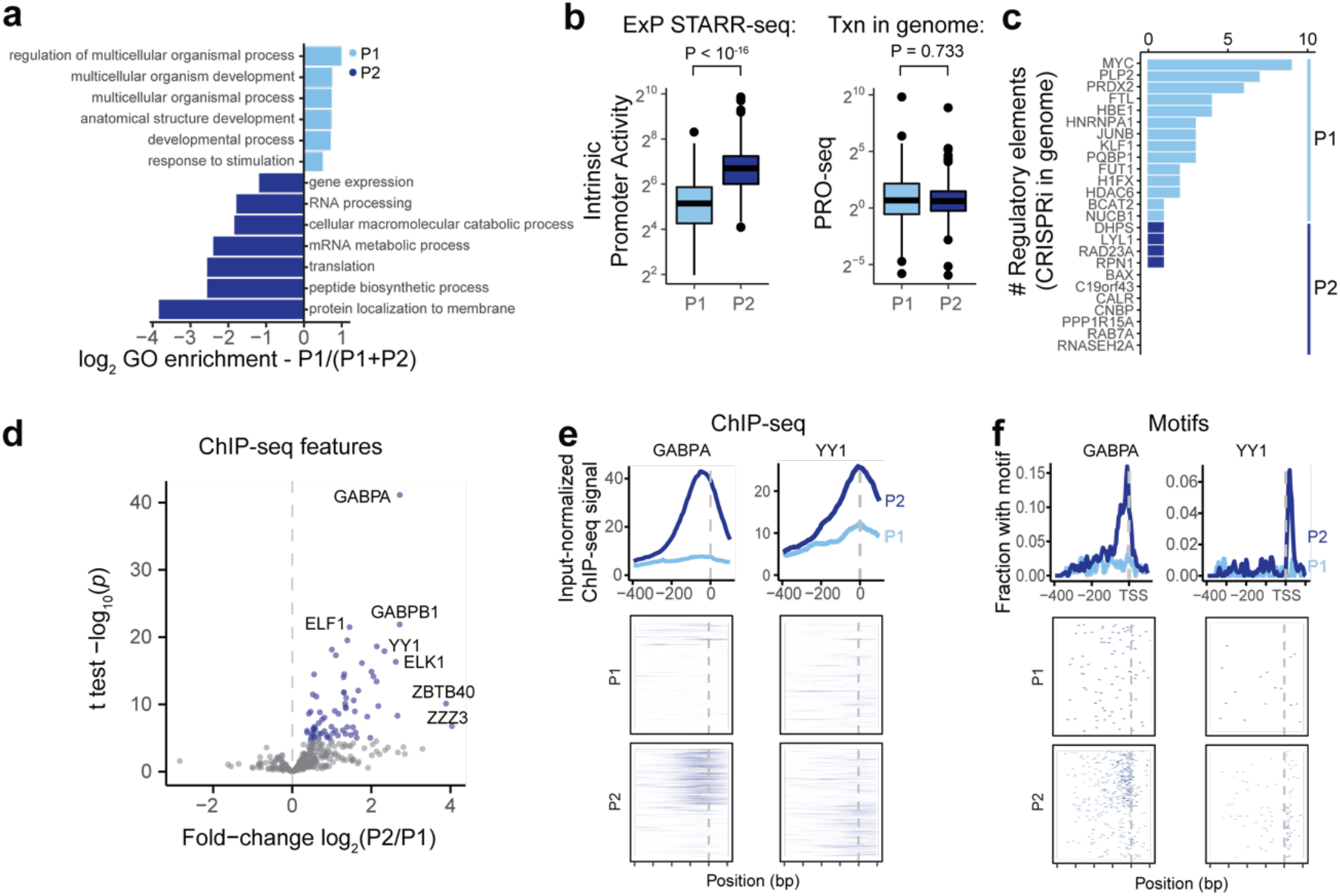
Promoter classes correspond to enhancer-responsive versus constitutive genes. **a.** Gene ontology log_2_-enrichment for P1 promoters using P1 and P2 promoters as a background set. **b.** Intrinsic promoter activity for P1 vs P2 promoters (ExP STARR-seq) and genomic transcription level of genes corresponding to P1 vs P2 promoters (PRO-seq reads per kilobase per million in gene bodies). **c.** Number of activating genomic regulatory elements identified in comprehensive CRISPRi screens for genes corresponding to P1 promoters (n=14) and P2 promoters (n=11)^22^. **d.** Volcano plot comparing ChIP-seq and other biochemical features for P2 versus P1 promoters (see **Table S6**). X-axis: ratio of average signal at P2 versus P1 promoters. Blue points: features with significantly higher signal at P2 promoters; no features have significantly higher signal at P1 promoters. **e.** ChIP-seq signal for GABPA and YY1 in K562 cells at P1 and P2 promoters in the genome, aligned by TSS (see Methods). Top: average ChIP signal (normalized to input) +/- 95% c.i. Bottom: signal at individual genomic promoters. **f.** Motif occurrences for GABPA and YY1 in P1 and P2 promoters, aligned by TSS.

P1 promoters had on average 3.2-fold weaker intrinsic promoter activity than P2 promoters, as measured by ExP-STARR-seq (P < 10^-16^, Mann-Whitney *U*-test; Fig. 4b), but showed similar levels of transcription in their native genomic locations, as measured by PRO-seq in the gene body (*P* = 0.733, Mann-Whitney *U*-test; Fig. 4b). This suggests that P1 promoters may be more dependent on genomic context for their level of transcription in the genome.

Genes corresponding to P1 promoters had more genomic regulatory elements in CRISPR experiments. In data from previous studies, in which CRISPRi was used to perturb every DNase-accessible element near selected promoters, the 14 genes corresponding to P1 promoters had an average of 3.6 (median: 3) distal enhancers in CRISPR experiments, whereas the 11 genes corresponding to P2 promoters had only 0.36 (median: 0, Fig. 4c), despite having similar numbers of nearby accessible elements (Fig. S6a). Distal enhancers for P1 genes in the genome also had stronger effect sizes (*P* = 0.0071, *t-*test, Fig. S6b).

Together, these observations suggest that P1 promoter sequences correspond to context-specific genes and depend more on distal enhancers for their transcriptional activation both in ExP STARR-seq and in the genome, whereas P2 promoter sequences correspond to constitutively expressed genes that are relatively less sensitive to distal enhancers in both contexts.

### TFs positioned at TSS distinguish constitutive from responsive promoters

We next sought to identify sequence and chromatin features that distinguish P1 (“responsive”) from P2 (“constitutive”) promoters.

We considered canonical core promoter motifs, which have been observed to differ between various subsets of promoters^29–33^, but did not find strong relationships. P1 and P2 promoter sequences had similar frequencies of the canonical ‘CA’ Initiator dinucleotide at the TSS (40.1% vs 35.3%, Fig. S6c), and corresponded to genes with similar patterns of dispersed versus focused TSSs in the genome (Fig. S6d). Consistent with previous studies comparing features of housekeeping versus other gene promoters^29–33^, P2 promoters had a slightly higher frequency of CpG dinucleotides (median 0.90 vs 0.81 normalized CpG content for P2 and P1 promoters, Fig. S6e), and P1 promoters had a 2-fold higher frequency of TATA box sequences upstream of the TSS (12.5% vs 6.1%), although only a small proportion of promoters contained this motif (Fig. S6c).

Accordingly, we explored which other sequence features or TF binding measurements distinguished P2 constitutive from P1 responsive promoters. We examined 3,206 other features (including ChIP-seq measurements, TF motif predictions, and other features), and identified striking differences in the frequencies of certain transcription factor binding sites and motifs (Fig. 4d, Fig. S6f, **Table S7**, see Methods). The most significantly enriched features included ChIP-seq signal for ETS family factors (GABPA, ELK1, ELF1), YY1, HCFC1, NR2C1, and C11orf30 / EMSY (Fig. 4d, Fig. S7). For example, two of the top factors (GABPA and YY1) together showed strong binding to a total of 64% of P2 promoters in the genome: 50% of P2 promoters showed strong GABPA binding (vs 8% of P1 promoters; *P* = 9.9 x 10^-22^, BH-corrected Fisher’s exact test), and 29% of P2 promoters showed strong YY1 binding (vs 5% of P1 promoters, *P* = 9.4 x 10^-9^, BH-corrected Fisher’s exact test) (Fig. 4e). Notably, the sequence motifs for these factors showed positional preferences consistent with a function in regulating transcription initiation: the motif for GABPA was typically located 0-20 nucleotides upstream of the TSS (mode: –10), and for YY1 was often positioned at either +18 bp (both strands) or +2 bp (negative strand) from the TSS (Fig. 4f, Fig. S6g). Consistent with these factors playing a functional role, previous studies have found that adding GABPA or YY1 motifs to promoters increases gene expression in various reporter assays and cell types^34–37^.

Together, these analyses suggest that P2 promoters can best be distinguished from P1 promoters by the presence of certain transcription factors including GABPA and YY1, rather than canonical core promoter motifs.

### P2 constitutive promoters contain ‘built-in’ enhancer sequences

We considered how transcription factors such as GABPA and YY1 might contribute to the reduced enhancer responsiveness of P2 versus P1 promoters. Interestingly, we noticed that these same factors showed strong binding in the genome not only at P2 promoters (Fig. 4e,f), but also at some E1 enhancers (Fig. S5a, Fig. S7b). For example, 3 of the genomic enhancers for *HBE1* (all classified as E1 in ExP STARR-seq) contained GABPA sequence motifs and showed strong GABPA binding by ChIP-seq, whereas the genomic promoter of *HBE1* (classified as P1) lacked these features (Fig. 5a).

**Fig. 5.**
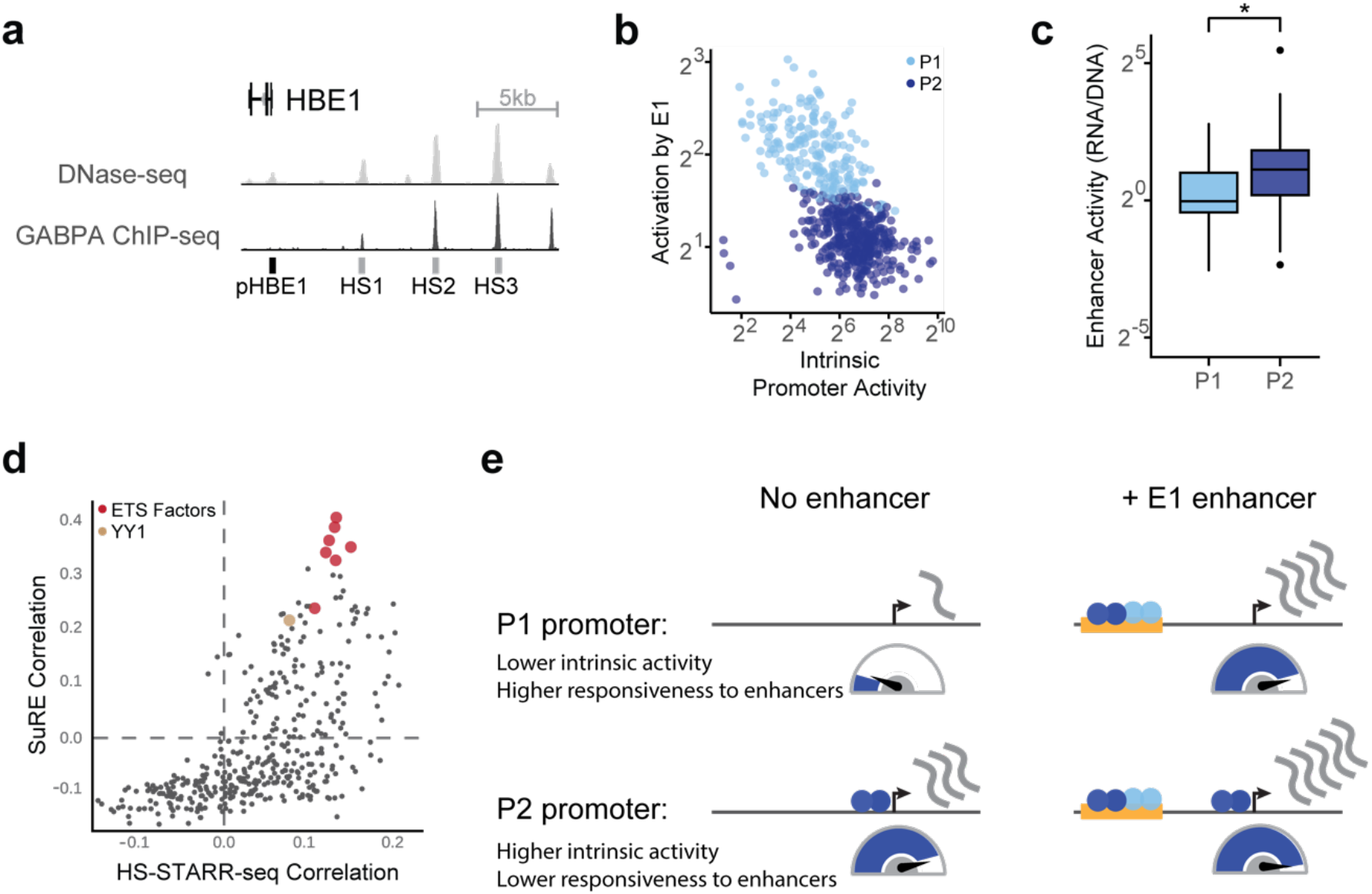
P2 constitutive promoters contain built-in enhancer sequences. **a.** DNase-seq and GABPA ChIP-seq binding at the HBE1 promoter (pHBE1) and HS1-HS3 enhancers. **b.** Correlation between intrinsic promoter activity and responsiveness of promoters to E1 enhancers (average activation by E1 sequences, expressions vs. random genomic controls). Each point is one promoter. **c.** Average enhancer activity in HS-STARR-seq (RNA/DNA) of P1 and P2 promoters. \**P* = 1.14 x 10^-16^, *t*-test. **d.** For each of 400 sequence motifs that appeared in at least 5% of HS-STARR-seq fragments, correlation (Pearson *R*) of motif occurrence with intrinsic promoter activity (SuRE signal, y-axis) and with intrinsic enhancer activity (HS-STARR-seq signal among fragments not overlapping TSS, x-axis). **e.** A model for enhancer-promoter compatibility. Enhancers multiplicatively scale the RNA output of promoters. P2 constitutive promoters contain built-in activating sequence motifs that both increase intrinsic promoter activity and reduce responsiveness to distal enhancers.

These observations suggested that P2 promoters may have reduced responsiveness to E1 enhancers because they contain some of the same motifs, potentially saturating some step in transcription. Accordingly, we explored the hypothesis that P2 promoters contain ‘built-in’ E1 enhancer sequences that would increase promoter activity and decrease responsiveness to distal E1 enhancers.

Consistent with this hypothesis, we found that (i) across all promoters, responsiveness to E1 enhancers was inversely correlated with intrinsic promoter activity, in a way that appeared to saturate; (ii) P2 promoters had stronger enhancer activity than P1 promoters; and (iii) nearly all of the TF motifs enriched in P2 promoters were predictive of both promoter activity and enhancer activity:

We first compared intrinsic promoter activity with responsiveness to E1 enhancers, and found that they were correlated both when considering all promoters in ExP STARR-seq (Pearson *R* = –0.62, log_2_ space; Fig. 5b) and when considering only P1 promoters (*R* = –0.51). For example, comparing P1 promoters at opposite extremes, the *RAD23A* promoter (P2) had 11.8-fold higher intrinsic promoter activity compared to the *HBE1* promoter (P1), and was 2.1-fold less sensitive to E1 enhancers. As promoter activity increased, responsiveness to E1 enhancers decreased rapidly (for example, from ∼9-fold average activation by E1 enhancers for the *SNAI3* P1 promoter) and appeared to saturate at ∼3-fold for most P2 promoter sequences (Fig. S8a).

We next tested whether P2 promoters had stronger intrinsic enhancer activity. To do so, we generated a second STARR-seq dataset in which we measured the enhancer activity of >8.9 million sequences derived from DNase-accessible elements and promoters (by hybrid selection (HS)-STARR-seq, see Methods, Fig. S8b-d). In this dataset, many promoter elements tested in ExP STARR-seq (along with thousands of other accessible elements) were densely tiled (an average of ∼11 fragments each covering at least 90% of the promoters tested in the ExP assay), allowing us to test the enhancer activity of entire P1 and P2 promoter sequences. P2 promoters indeed showed ∼2-fold higher intrinsic enhancer activity than P1 promoters in HS-STARR-seq (*P* = 1.14 x 10^-16^, *t*-test, Fig. 5c), supporting a model where these promoters contain built-in enhancers.

Finally, we examined whether the sequence motifs enriched in P2 promoters contribute to both enhancer activity and promoter activity. To do so, we examined data on enhancer activity from HS-STARR-seq along with another previous experiment that measured promoter activity for millions of random genomic fragments in K562 cells (SuRE^38^). 16 of the 17 motifs enriched in P2 promoters, including motifs for GABPA and YY1, were positively correlated with both enhancer activity and promoter activity (Fig. 5d, **Table S7**, see Methods).

Together, these observations suggest a model for promoter sequence organization (Fig. 5e). P2 promoters encode binding motifs for activating factors, including GABPA and YY1, that act as ‘built-in’ enhancers for the promoter. This not only increases the autonomous activity of the promoter, but also reduces its responsiveness to distal enhancers. P1 promoters, in contrast, appear to exclude these activating factors, creating a sensitivity to distal enhancers.

## Discussion

Since the discovery of the first enhancers forty years ago^1, 2^, many enhancer and promoter sequences have been combined and found to be compatible^3–9^. At the same time, studies of individual natural or synthetic core promoters have been found to have some degree of specificity when combined with various transcriptional cofactors or enhancer sequences^10–15^.

Here we develop and apply ExP STARR-seq to systematically quantify enhancer-promoter compatibility, and identify a simple rule for combining human enhancer and promoter activities. Enhancers are intrinsically compatible with many Pol II promoter sequences, and act multiplicatively to scale the RNA output of a promoter. As a result, independent control of intrinsic enhancer activity and intrinsic promoter activity can create significant variation in RNA expression: in our data, promoter activity and enhancer activity each vary over 3-4 orders of magnitude, and their multiplicative combination leads to >4-million-fold variation in STARR-seq expression. This finding of broad compatibility appears to be consistent with recent studies using reporters integrated into the genome, which found that human core promoters or enhancers are similarly scaled when they are inserted into different genomic loci^39, 40^. This is also consistent with our previous finding that the effects of enhancers on nearby genes in the genome can be predicted with good accuracy using a model based only on genomic measurements of enhancer activity and distance-based 3D contacts, assuming no intrinsic enhancer-promoter specificity^22^. While there may be circumstances where promoters are responsive only to certain cofactors or enhancer sequences^10–15^, our observations indicate that biochemical specificity is not the dominant factor controlling the activity levels of human enhancers and promoters.

Superimposed on this multiplicative function, we identify two classes of enhancers and promoters that show subtle preferences in activation. One class of promoters, corresponding largely to constitutively expressed (housekeeping) genes, is less responsive to distal enhancers both in ExP STARR-seq and in the genome, while the second class of promoters, corresponding to cell-type- or context-specific genes, is more responsive. Previous studies have identified numerous differences in sequence content and motifs between the promoters of housekeeping and context-specific genes^29–33^. We find that these promoters indeed show intrinsic differences in their levels of activity and responsiveness to enhancers. Interestingly, this pattern of promoter responsiveness also can be predicted by a simple logic: P2 promoters contain built-in activating sequences that increase both enhancer and promoter activity, which appears to reduce their responsiveness to distal enhancers. This model for human promoters appears to differ qualitatively from previous studies in *Drosophila*, which found that the promoters of housekeeping and developmentally regulated genes can both be highly responsive, but to distinct sets of enhancer sequences and cofactors^16, 18^. We note that one methodological difference is that, whereas these previous studies focused on minimal core promoter sequences (100-138bp total), here we included more sequence context upstream of the TSS (264 bp total).

A remaining challenge will be to link the sequences that control enhancer and promoter activities with effects on particular biochemical steps in transcription. In this regard, we find that GABPA and YY1 bind both to constitutive promoters and to distal enhancers, and are associated with increased enhancer activity, increased promoter activity, and reduced promoter responsiveness to distal enhancers. This suggests that distal enhancers may act, in part, on a particular rate-limiting step in transcription that can be saturated by inclusion of built-in activating sequences in a gene promoter. Indeed, a previous study found that adding GABPA and YY1 motifs to several promoters led to an increase in RNA expression that saturates at 2 or 5 copies of the motif, respectively.^34^ Given the preferred positions of these motifs within 20 bp of the TSS — as well as previous findings that these proteins physically interact with general transcription factors^41, 42^ and/or influence transcriptional initiation and TSS selection^36, 43–45^ — such a rate-limiting step might involve assembly of the preinitiation complex. In addition to this step, our data are consistent with a model in which enhancers and promoters control additional steps in transcription that combine multiplicatively and do not saturate in the dynamic range of our assay. Examples of such processes that could combine multiplicatively include control of burst frequency and burst size^46^. Further work will be required to investigate these possibilities.

Together, our findings support a simple logic for human enhancer-promoter compatibility, and will propel efforts to model gene expression, map the effects of human genetic variation, and design regulatory sequences for gene therapies.

## Acknowledgements

This work was supported by an NHGRI Genomic Innovator Award (R35HG011324 to J.M.E.); Gordon and Betty Moore and the BASE Research Initiative at the Lucile Packard Children’s Hospital at Stanford University (J.M.E.); an NIH Pathway to Independence Award (K99HG009917 and R00HG009917 to J.M.E.); the Harvard Society of Fellows (J.M.E.); the Broad Institute (E.S.L.); an AΩA Carolyn L. Kuckein Student Research Fellowship (D.T.B.); and by the National Institute of General Medical Sciences (T32GM007753, L.S.). We thank C. Vockley, V. Subramanian, and members of the Engreitz and Lander labs for discussions and technical assistance.

## Author Contributions

D.T.B., C.P.F., T.R.J., and J.M.E. developed the ExP STARR-seq assay. D.T.B., M.K., and T.H.N. performed experiments. D.T.B., T.R.J., V.L., L.S., H.Y.K., J.N., S.R.G., and J.M.E. analyzed data. E.S.L. and J.M.E. supervised the work. All authors contributed to writing the manuscript.

## Data Availability

Raw and processed data for ExP STARR-seq and HS STARR-seq can be found in NCBI GEO under accession number GSE184426. Luciferase data can be found in Supplementary Table S3.

## Tables

**S1.** Gene promoters used in ExP STARR-seq

**S2.** Candidate enhancers used in ExP STARR-seq

**S3.** ExP-Luciferase elements and data

**S4.** Biochemical feature enrichment in E1 vs. E2 enhancers

**S5.** Transcription factor motif enrichment in E1 vs. E2 enhancers

**S6.** Biochemical feature enrichment in P1 vs. P2 promoters

**S7.** Transcription factor motif enrichment in P1 vs. P2 promoters

**S8.** Primer and oligo sequences

**S9.** ENCODE datasets used to annotate ExP enhancers and promoters

**S10**. Enhancer hybrid selection probe sequences

**S11**. Promoter hybrid selection probe sequences

## Supplementary Figures

**Fig. S1.**
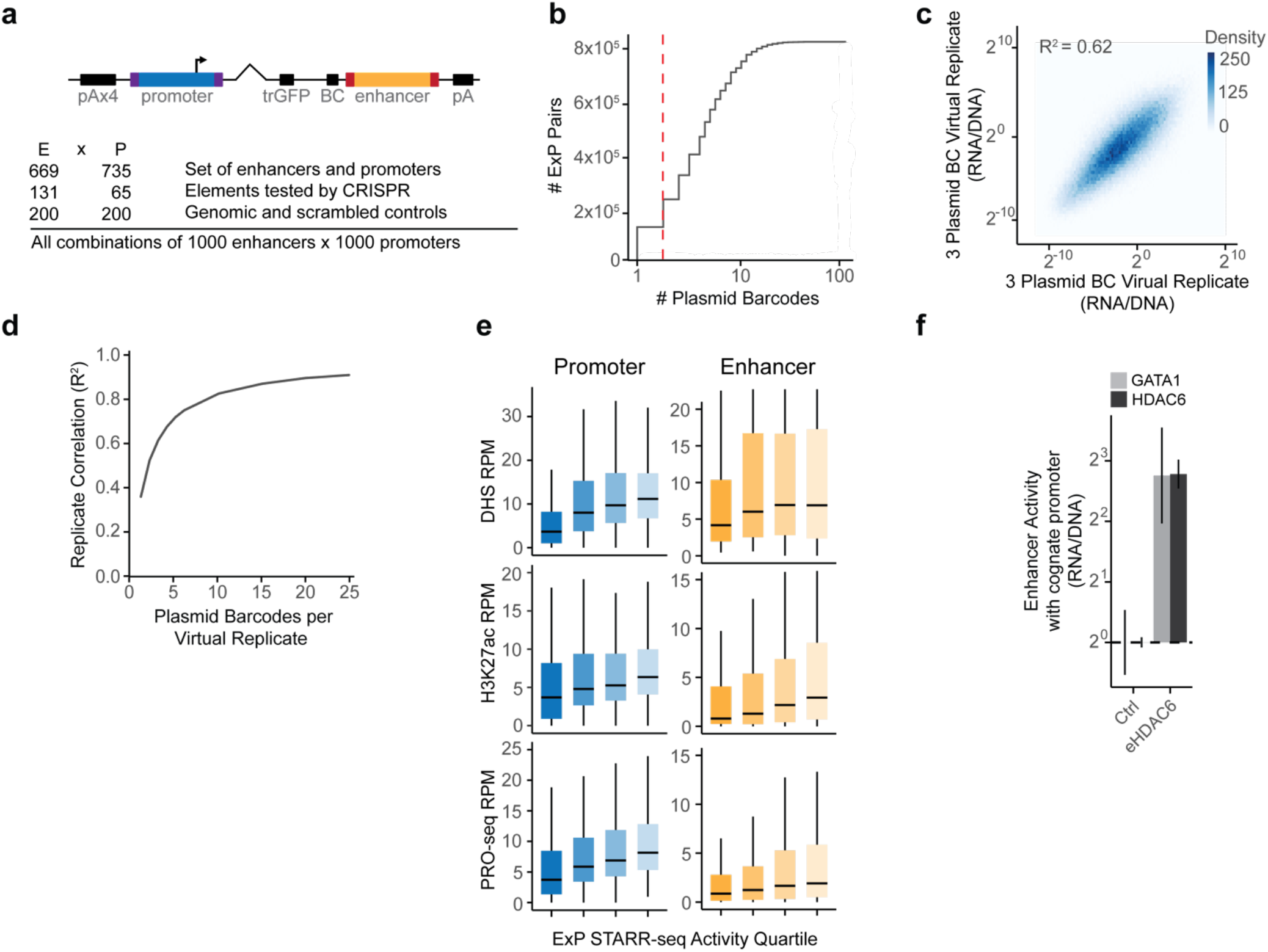
Design and reproducibility of ExP STARR-seq. **a.** ExP STARR-seq reporter construct (pA = polyadenylation signal; purple = promoter sequencing adaptors; angled = spliced sequence; trGFP = truncated GFP open reading frame; BC = 16bp N-mer plasmid barcode; red = enhancer sequencing adaptors) and 1000×1000 K562 library contents. **b.** Distribution of plasmid barcodes per enhancer-promoter pair, red dotted-line is threshold of two plasmid barcodes. **c.** Correlation between virtual replicates, formed by sampling two nonoverlapping groups of three plasmid barcodes from pairs with at least 6 barcodes, and averaging log_2_(RNA/DNA) within groups. **d.** Correlation between virtual replicates as in (**c**) for increasing numbers of plasmid barcodes per pair in virtual replicates. **e.** DNase-seq, H3K27ac ChIP-seq, and PRO-seq (RPM) by increasing quartile of autonomous promoter activity and average enhancer activity in ExP STARR-seq. Box: median and interquartile range (IQR). Whiskers: +/- 1.5 x IQR. **f.** Activation in ExP STARR-seq (expression versus genomic controls in distal position) of GATA1 and HDAC6 promoters by eHDAC6 (chrX:48641342-48641606). Ctrl = activity of promoters with random genomic controls in enhancer position. Error bars: 95% CI across plasmid barcodes. n = 7 (GATA1-ctrl), 381 (HDAC6-ctrl), 4 (eHDAC6-GATA1), 37 (eHDAC6-HDAC6).

**Fig. S2.**
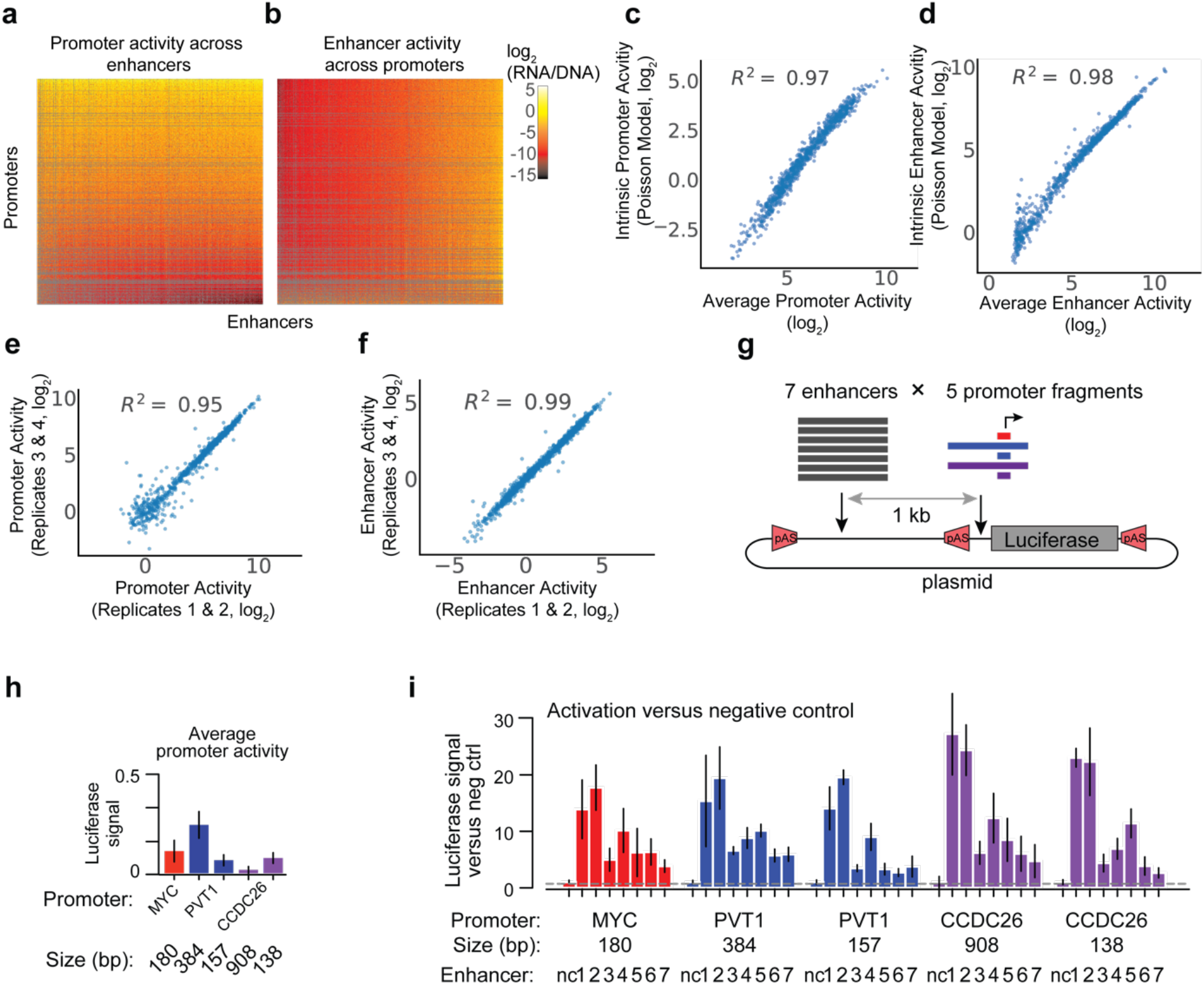
Comparison of methods of estimating enhancer and promoter activities and validation of multiplicative model using luciferase assays. **a-b.** Heatmap of promoter activity (**a,** expression divided by intrinsic enhancer activity) or enhancer activity (**b,** expression divided by instrinsic promoter activity) across all pairs of promoter (vertical) and enhancer sequences (horizontal). Axes are sorted by intrinsic promoter and enhancer activities, as in Fig. 2j. Grey: missing data. **c-d.** Correlation between two estimates of promoter (**c**) and enhancer (**d**) activities. One method (“average activity”, x-axis) estimates activity calculated by averaging across elements, and the other method (“intrinsic activity”, y-axis) estimates activity by using coefficients estimated by a Poisson count model (see Methods). **e-f.** Correlation of intrinsic promoter (**e**) and enhancer (**f**) activity estimates from Poisson model using data from separate replicate experiments. **g.** ExP luciferase reporter construct. Seven enhancer fragments, with flanking polyadenylation signals, were cloned upstream of five promoter fragments and measured via the dual luciferase assay. **h.** Autonomous promoter activity of ExP luciferase (average luciferase signal of promoter with negative control) for 5 promoter sequences derived from 3 genes (*MYC*, *PVT1*, *CCDC26*). Error bars are 95% CI from three biological replicates. **i.** Enhancer activation (luciferase signal versus negative control sequence in the enhancer position) of seven enhancers across five promoter fragments. Error bars are 95% CI from three biological replicates.

**Fig. S3.**
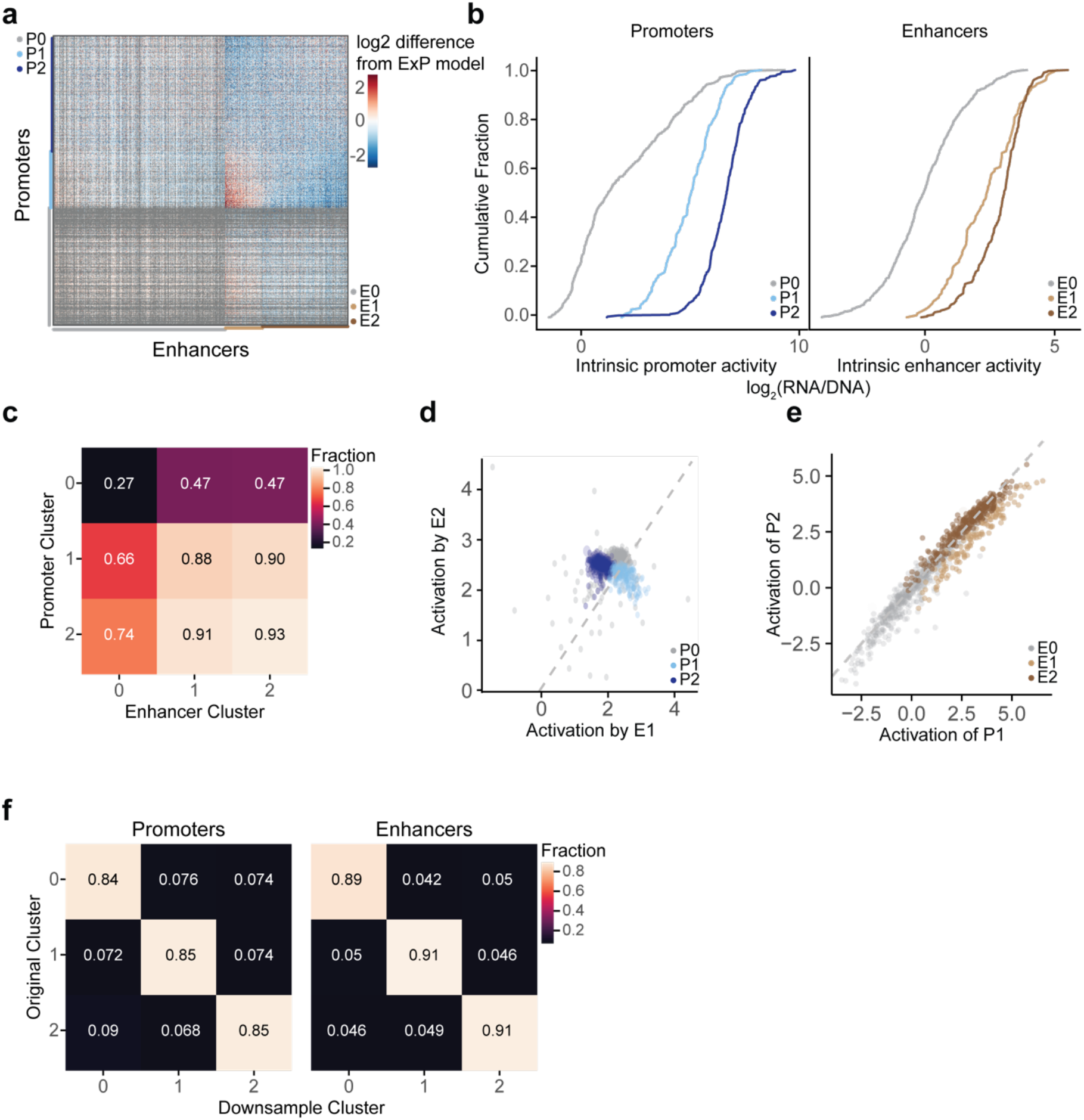
Enhancer and promoter cluster identification and reproducibility. **a.** Heatmap of deviations in enhancer-promoter STARR-seq expression from a multiplicative enhancer-promoter model (color scale: fold-difference between observed expression versus expression predicted by multiplicative model; gray: missing data). Same as Fig 3a, except including clusters with weak sequences and missing data (E0 and P0). Vertical axis: promoter sequences grouped by class and sorted by responsiveness to E1 vs. E2; horizontal axis: enhancer sequences grouped by class and sorted by activation of P1 vs. P2. **b.** Distribution of intrinsic enhancer and promoter activity (expression versus genomic controls) by cluster. **c.** Fraction of enhancer-promoter pairs observed in ExP STARR-seq dataset (>= 2 plasmid barcodes) by cluster. **d.** Correlation of average promoter activation (expression versus genomic controls in enhancer position) by E2 versus E1 enhancer sequences. Each point is one promoter sequence. Same as Fig. 3c, except including P0 promoter sequences. **e.** Correlation of average activation of P2 versus P1 promoters. Each point is one enhancer sequence. Same as Fig. 3d, except including E0 enhancer sequences. **f.** Robustness of enhancer and promoter cluster assignments to downsampling of enhancer and promoter sequences. Clustering was repeated in 100 random downsamplings to 25% of promoter sequences and 25% of enhancer sequences (6.25% of original matrix). Heatmap: Average fraction overlap between cluster assignments from the full and downsampled matrices.

**Fig. S4.**
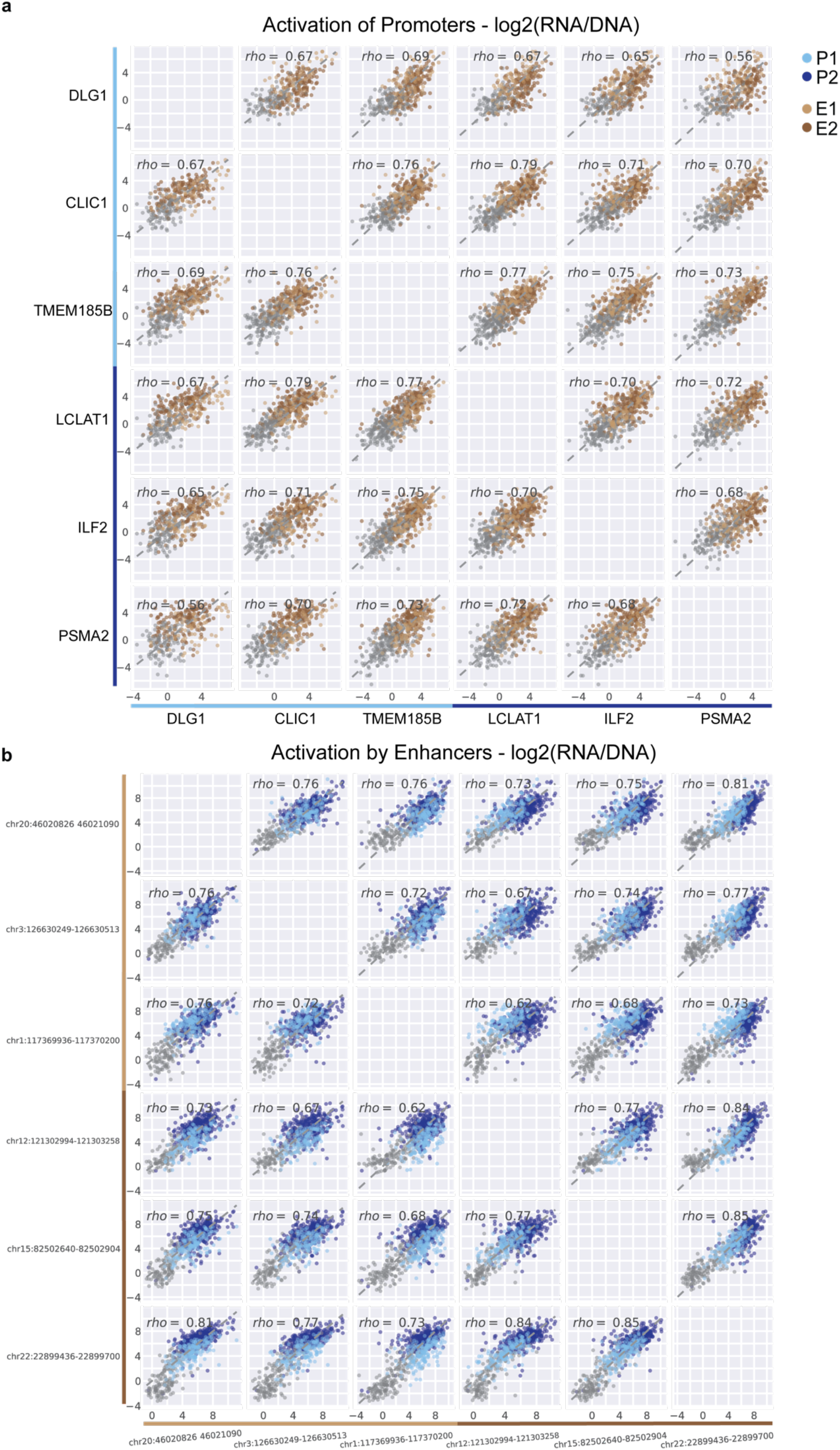
Classes of enhancer and promoter sequences show distinct patterns of activation and responsiveness. **a.** For 6 representative promoter sequences (3 P2 and 3 P1 sequences), the pairwise correlation of activation by enhancers (expression versus genomic controls in enhancer position, averaged across plasmid barcodes). Each point is one enhancer sequence. **b.** For 6 representative enhancer sequences (3 E1 and 3 E2 sequences), the pairwise correlation of promoter activation (expression versus genomic controls in promoter position, averaged across plasmid barcodes). Each point is one promoter sequence.

**Fig. S5.**
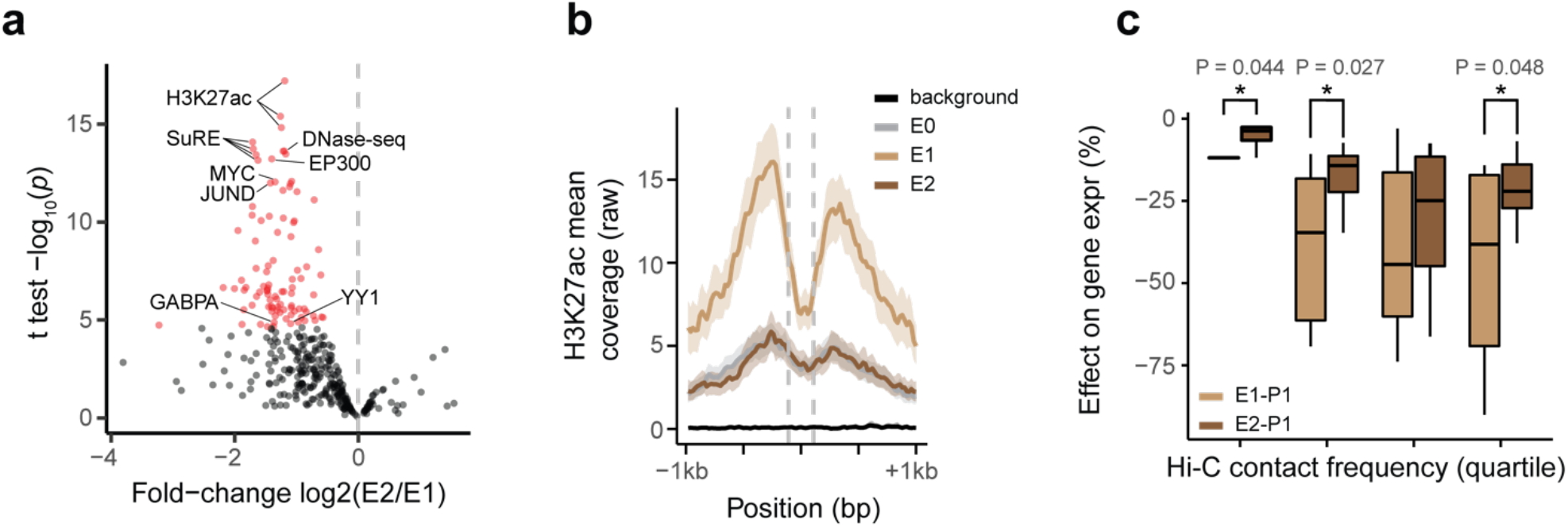
Classes of enhancer sequences correspond to strong and weak genomic enhancers. **a.** Volcano plot comparing ChIP-seq and other genomic features for E2 versus E1 enhancer sequences (see **Table S4**). X-axis: ratio of average signal at P2 versus P1 promoters. Red dots: features with significantly higher signal at E1; no features have significantly higher signal at E2 enhancer sequences. **b.** Mean H3K27ac ChIP-seq coverage of genomic elements corresponding to E0, E1, E2, or genomic control enhancer sequences (+/- 95% CI), aligned by DHS peak summit. Dotted lines mark bounds of the enhancer sequences used in ExP STARR-seq. **c.** % effect of genomic elements corresponding to E1 vs. E2 enhancer sequences on expression of genes corresponding to P1 promoters in CRISPRi screens, separated by quartiles of 3D contact frequency measured by Hi-C (0.39-11.9, 11.9-23.9, 23.9-58.3, 58.3-100). \**P* < 0.05, two-sample *t*-test.

**Fig. S6.**
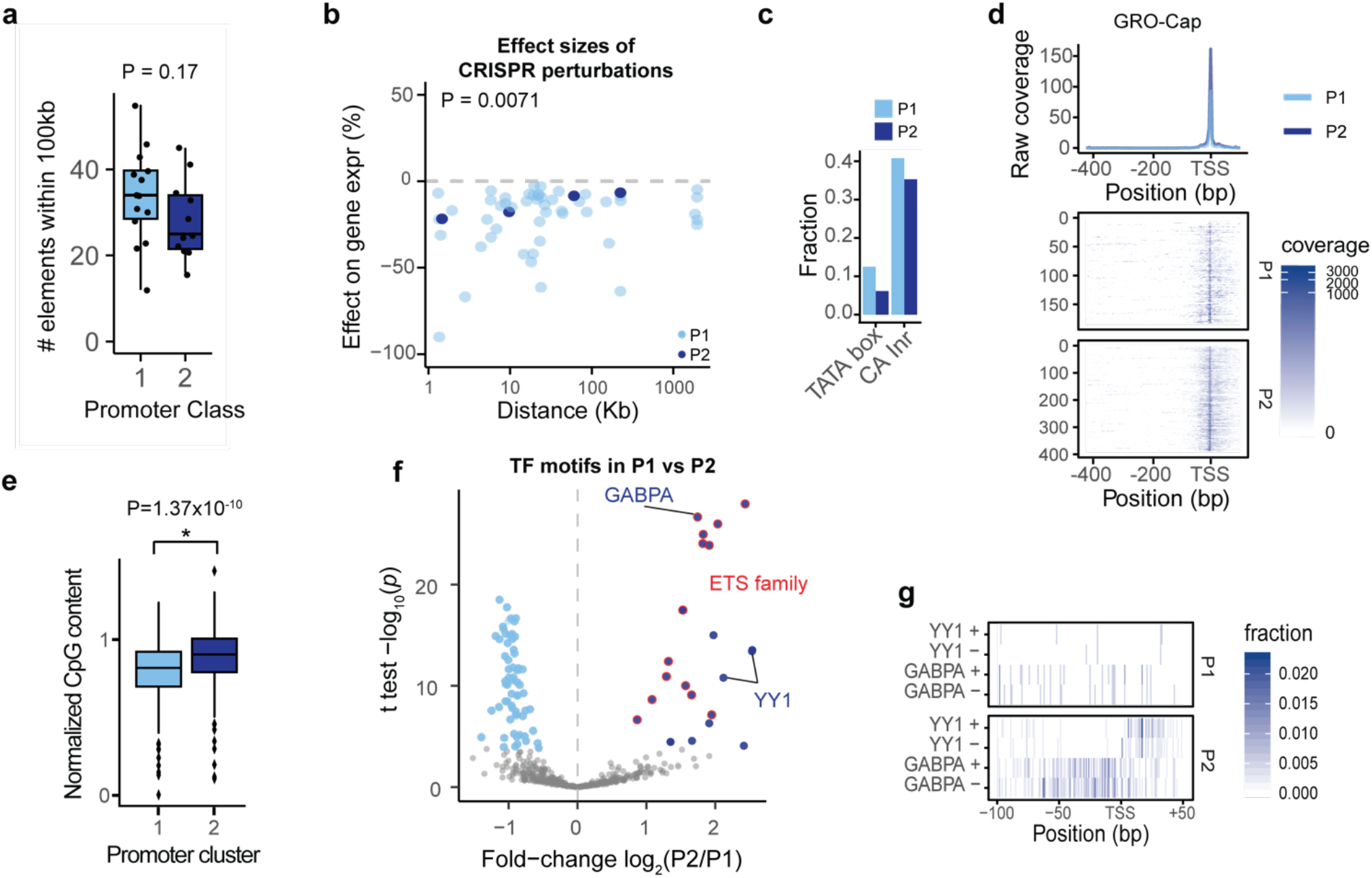
Properties of promoter classes. **a.** Number of nearby accessible elements (within 100 Kb of the gene promoter, considering top 150,000 DNase peaks in K562 cells as used in the ABC model^22^) for the 14 genes corresponding to P1 promoters and 11 genes corresponding to P2 promoters with comprehensive CRISPR tiling data. *P* = 0.17, Mann-Whitney U test. **b. %** Effect of CRISPRi perturbations to genomic regulatory elements on genes corresponding to P1 vs. P2 promoters. *P* = 0.0071, *t-*test. **c.** Fraction of promoter sequences containing TATA or CA initiator core promoter motifs. **d.** GRO-Cap coverage of genomic promoters aligned by TSS. Top: Mean coverage of genomic promoters corresponding to P1 vs. P2 classes. Bottom: Coverage across all individual promoters. **e.** Normalized CpG-content of P1 and P2 promoter sequences, calculated as the ratio of observed to expected CpG = (CpG fraction) / ((GC content)^2^ / 2). **f.** Volcano plot comparing frequency of transcription factor motifs in P2 versus P1 promoter sequences (see **Table S7**). X-axis: ratio of average motif counts in P2 versus P1 promoter sequences. Light blue and dark blue dots: Motifs significantly more frequent in P1 or P2 promoter sequences, respectively. Red outline: significant motifs for ETS family transcription factors. **g.** Fraction of P2 promoter sequences with YY1 and GABPA binding motifs by nucleotide position, aligned by TSS and separated by strand (see Methods).

**Fig. S7.**
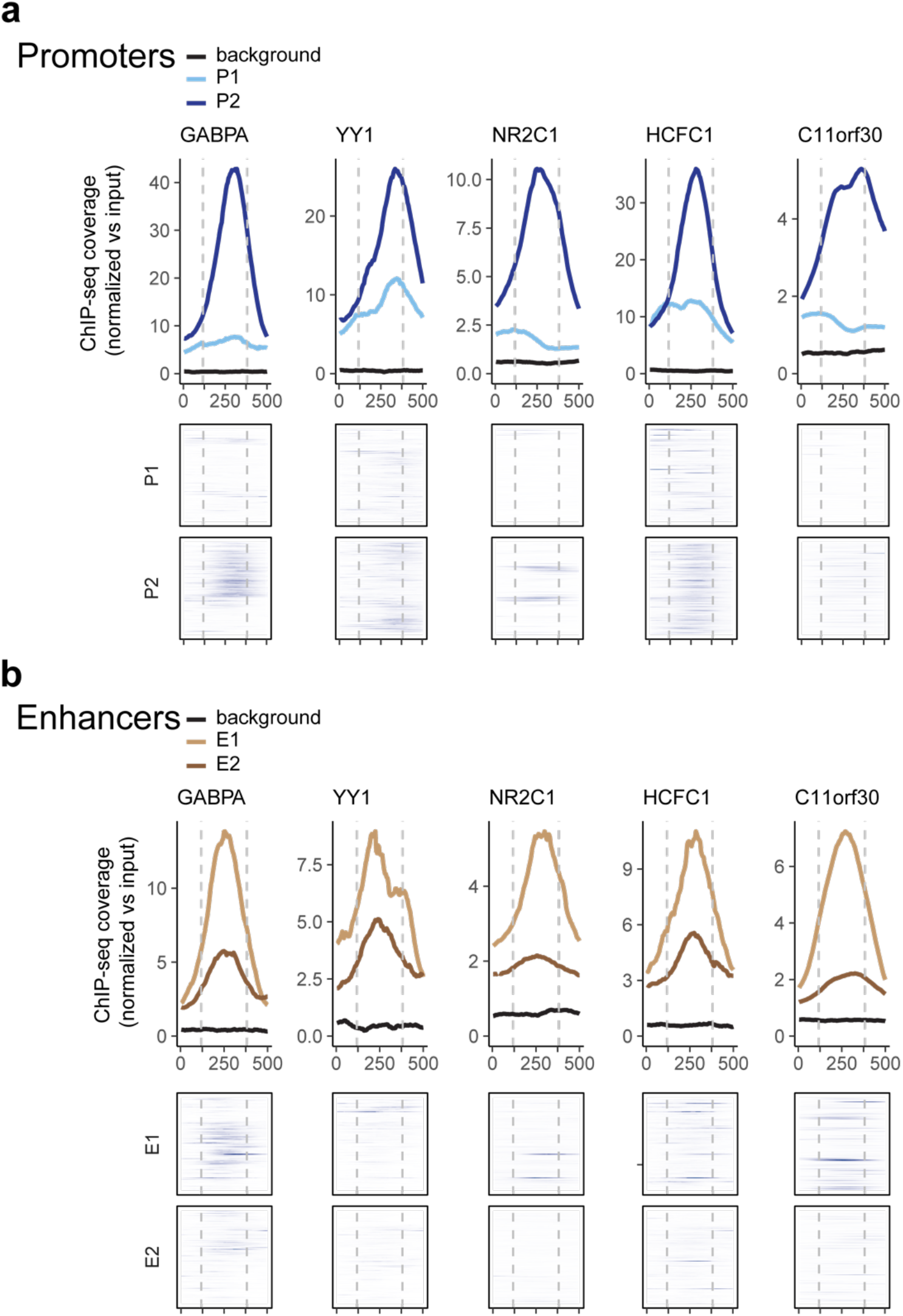
Transcription factors enriched at P2 promoters are also enriched at E1 enhancers. **a.** ChIP-seq signal for 5 transcription factors in K562 cells at P1 and P2 promoters in the genome, aligned by boundaries of the 264-bp ExP STARR-seq promoter sequence (see Methods). Top: average ChIP-seq signal normalized to input. Bottom: signal at individual genomic promoters. Black line: average for random genomic control sequences. **b.** ChIP-seq signal at E1 and E2 enhancers in the genome. Black line: average for random genomic control sequences.

**Fig. S8.**
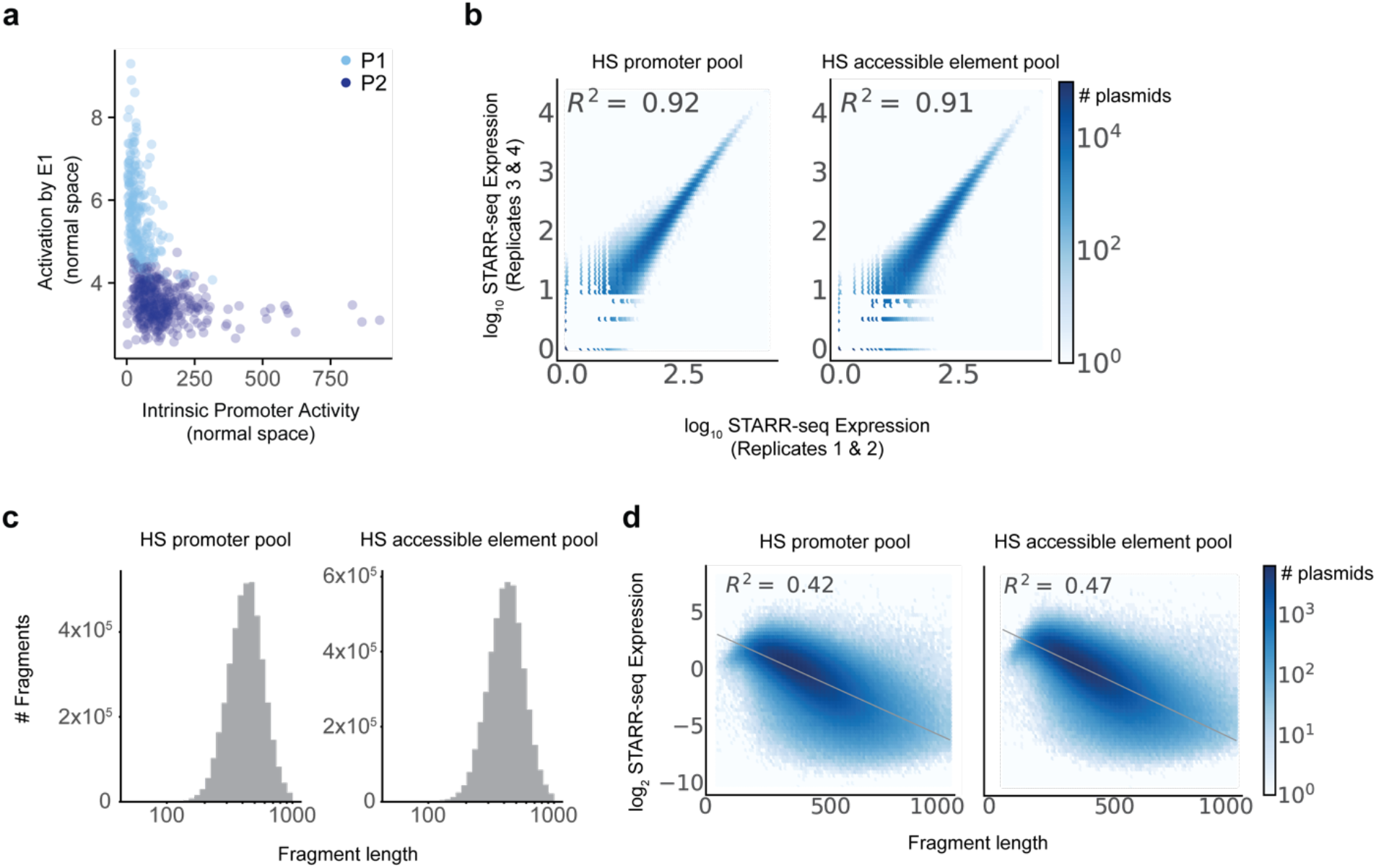
Responsiveness to E1 enhancers versus intrinsic promoter activity, and metrics for hybrid-selection STARR-seq experiments. **a.** Correlation between intrinsic promoter activity and responsiveness of promoters to E1 enhancers (average activation by E1 sequences, expressions vs. random genomic controls). Each point is one promoter. Same as Fig. 5b, but in normal scale instead of log_2_ scale. **b.** Correlation of HS-STARR-seq expression between biological replicate experiments for promoter and accessible element pools, calculated for individual elements with unique plasmid barcodes. Axes represent the average STARR-seq expression (RNA/DNA, log_10_ scale) of two biological replicates. Density: number of plasmids. **c.** Fragment length distribution in HS-STARR-seq in promoter and accessible element pools, of fragments with at least 25 DNA counts. **d.** STARR-seq expression (y-axis) and fragment length (x-axis) relationship in HS-STARR-seq. Density: number of plasmids.

## Methods

### Genome build

All analyses and coordinates are reported using human genome reference hg19.

### Design of ExP STARR-seq

We designed ExP STARR-seq to systematically measure the intrinsic, sequence-encoded compatibility or specificity of many pairs of human enhancer and promoter sequences. The key design features we considered when developing this assay were the ability to measure the activity of individual enhancer-promoter sequence combinations, to precisely quantify the expression of each enhancer-promoter pair, and to test hundreds of thousands of combinations in order to identify patterns of compatibility or specificity across a large number of human sequences.

Accordingly, we designed a new variant of the STARR-seq high-throughput plasmid reporter assay called enhancer x promoter (ExP) STARR-seq. In both STARR-seq and ExP STARR-seq, enhancer sequences are cloned downstream of a promoter, transfected into cells, and transcribed to produce a reporter mRNA transcript, which is then sequenced to quantify the relative expression levels of plasmids containing different enhancer sequences^4^. In ExP STARR-seq, we modified the cloning and RNA sequencing strategy to enable testing different enhancer sequences in combination with different promoter sequences.

To clone combinations of enhancer and promoter sequences into a reporter plasmid, we synthesized 264-bp enhancer and promoter sequences in an oligo pool format, PCR amplified enhancer and promoter sequences separately, and inserted them into the hSTARR-seq_SCP1 vector_blocking 4 vector^8^ in the promoter position (replacing the original SCP1 promoter) or enhancer position in a single pooled cloning step using Gibson assembly to generate all pairwise combinations of chosen enhancer and promoter sequences (Fig. 1a, Fig. S1a). We chose this specific STARR-seq vector with 4 polyA sequences upstream of the promoter position because it was specifically designed in order to avoid spurious transcription initiation from the origin of replication^8^, which would interfere with the STARR-seq signal from the cloned enhancer-promoter pairs. This STARR-seq vector also includes 5’ and 3’ splice sites upstream of the enhancer that allows using a PCR primer targeting the splice junction to specifically amplify cDNA derived from the reporter mRNA while avoiding amplifying the plasmid DNA sequence.

To quantify the reporter mRNA transcripts and determine which enhancer-promoter pair they correspond to, we further adapted the cloning and RNA sequencing design. In the standard STARR-seq assay, the reporter mRNA contains the enhancer sequence but not the full promoter sequence, and therefore cannot determine from which promoter a given reporter mRNA is derived. Accordingly, in ExP STARR-seq we introduced a random 16-mer sequence located just upstream of the enhancer sequence that we use as a “plasmid barcode” to identify which enhancer reporter mRNAs are derived from which enhancer-promoter pairs (Fig. 1a). After cloning the plasmid pool, we map which plasmid barcodes correspond to which promoters by applying Illumina high-throughput sequencing to a PCR amplicon containing the promoter sequence and plasmid barcode. From this, we build a dictionary to look up, for a given reporter mRNA containing a plasmid barcode and enhancer sequence, which enhancer-promoter-plasmid barcode construct that mRNA is derived from.

Finally, we selected the number of constructed tested (∼1 million pairs of enhancer and promoter sequences cloned, with an average of 6.3 plasmid barcodes per pair) and sequencing depth (>1 billion reads per replicate) to enable highly precise measurements of expression for each enhancer-promoter pair. We obtained high reproducibility of enhancer-promoter expression levels between biological replicates (R^2^ = 0.92), allowing us to develop quantitative models of how enhancer and promoter activities combine.

Altogether, this approach enables precisely quantifying the expression levels of thousands of combinations of enhancer and promoter sequences.

### Selection of enhancer and promoter sequences for ExP STARR-seq

To explore the compatibility of human enhancers and promoters, we selected 1000 promoter and 1000 enhancer sequences, including sequences from the human genome spanning a range of expression or activity levels, and dinucleotide shuffled controls. Based on available lengths of oligonucleotide pool synthesis, each sequence was 264bp.

Promoters: We selected the 1000 promoter sequences to include:

- 65 genes whose enhancers have previously been studied in CRISPR experiments in K562 cells^22^
- 715 genes sampled to span a range of potential promoter activities, including the 200 most highly expressed genes in K562 cells, based on CAGE signal at their TSS^21^ and a random sample of 515 other expressed genes (>1 TPM in RNA-seq data^27^).
- 20 genes that are not expressed or lowly expressed in K562 cells (<1 TPM), and that are expressed in both GM12878 and HCT-116 cells (in the top 70% of genes by TPM based on RNA-seq^21^.
- 100 random genomic sequences as negative controls (+ strand)
- 100 dinucleotide shuffles of these random genomic sequences

For the selected genes, we synthesized a 264-bp sequence including approximately 244 bp upstream and 20 bp downstream of the TSS. Here, we defined the TSS as the center of the 10-bp window with the most CAGE 5’ read counts within 1 Kbp of a RefSeq TSS. For lowly expressed genes (which lack clear CAGE signal), we used the hg19 RefSeq-annotated TSS. For genes studied in Fulco *et al.* 2019, we adjusted the assigned 10bp TSS window by manual examination of the CAGE if necessary.

Enhancers: We selected the 1000 enhancer sequences to include:

- 131 elements previously studied with CRISPR^22^, including (i) all distal elements (i.e., >1 Kb from an annotated TSS) with significant effects in previous CRISPRi tiling screens (activating or repressive), (ii) all distal elements predicted by the Activity-by-Contact model to regulate one of the tested genes in K562 cells^22^, and (iii) two promoter elements for PVT1 that also act as enhancers for MYC^22^. We selected 264-bp regions centered on the overlapping DHS narrow peak. For the small number of CRISPR elements that did not overlap a narrow peak, we tiled the corresponding element with 264-bp windows overlapping by 50 bp.
- 200 DNase peaks with the strongest predicted enhancer activity, and 351 other DNase peaks sampled evenly across the range of predicted enhancer activity. Here, we considered all distal DHS peaks in K562 cells (DHS narrow peaks^22^) and calculated predicted enhancer activity as the geometric mean of DNase I hypersensitivity and H3K27ac ChIP-seq read counts in K562 cells in the ∼500-bp candidate enhancer regions used by the ABC model in Fulco et al. 2019^22^. Some candidate ABC elements in this set span more than one DHS peak, in which case we divided the predicted enhancer activity equally among each overlapping peak. We downloaded introns from the UCSC Genome Browser ‘refGene’ track (version 2017-06-24), and removed any peaks overlapping an annotated splice donor or acceptor site. We then selected 264-bp regions centered on the remaining DHS narrow peaks.
- 100 random genomic sequences as negative controls
- 100 dinucleotide shuffles of these random genomic sequences

All enhancer sequences were taken from the hg19 reference in the + strand direction.

### Library Cloning

We ordered 264bp sequences in an oligo array format from Twist Bioscience with separate pairs of 18bp adaptors (total length = 300bp) for enhancers (5’ = GCTAACTTCTACCCATGC, 3’ = GCAAGTTAAGTAGGTCGT) and promoters (5’ = TCATGTGGGACATCAAGC, 3’ = GCATAGTGAGTCCACCTT). We then PCR amplified enhancers and promoters separately from the same array using Q5 high-fidelity DNA polymerase (NEB M0492). We amplified enhancers in four 50uL PCR reactions (98°C for 30 seconds; 15 cycles of 98°C for 15 seconds, 61°C for 15 seconds, and 72°C for 20 seconds) using primers (forward: TAGATTGATCTAGAGCATGCANNNNNNNNNNNNNNNNGAGTACTGGTATGTTCAGCTAACTTCTACCCATGC, reverse: TCGAAGCGGCCGGCCGAATTCGTCATTCCATGGCATCTCACGACCTACTTAACTTGC) which add an additional 17bp on either side, a 16bp N-mer plasmid barcode upstream, and homology arms for Gibson assembly on either side of the enhancer sequence. We amplified promoters in four 50uL PCR reactions (98°C for 30 seconds; 4 cycles of 98°C for 15 seconds, 61°C for 15 seconds, 72°C for 20 seconds; 11 cycles of 98°C for 15 seconds and 72°C for 20 seconds) using primers (forward: CTCTGGCCTAACTGGCCGGTACGAGTGAGCTCTCGTTCATCATGTGGGACATCAAGC, reverse: CCCAGTGCCTCACGACCGGGCCTGGTAGCAAGCTTAGATAAGGTGGACTCACTATGC) which add an additional 17bp and homology arms for Gibson assembly on either side of the promoter sequence. We purified the PCR products using 0.8X volume of AMPure XP beads (Beckman Coulter, A63881) and pooled the reactions together while keeping enhancers and promoters separate.

We digested the human STARR-seq screening vector (hSTARR-seq_SCP1 vector_blocking 4, Addgene #99319) with both Thermo SgrDI and BshTI (AgeI) (replaced with enhancer sequence), then NEB KpnI and ApaI (replaced with promoter sequence), with purification using 0.8X volume AMPure XP after each digestion. We then recombined 500ng of this digestion (including ∼4.4kb of backbone vector and 250bp of filler sequence including a spliced region and truncated GFP ORF) with 150ng of both the purified enhancer and promoter products using Gibson assembly (NEB, E2611) for 1 hour at 50°C in a 40uL reaction and purified the reaction using 1X volume AMPure XP with 3 total ethanol washes.

We electroporated the assembled libraries into Lucigen Endura Electrocompetent cells (60242) using 0.1cm cuvettes (BioRad) using the Gene Pulser Xcell Microbial System (BioRad) (10 uF, 600 Ohms, 1800 Volts) following the manufacturer’s recommendations. We expanded the transformations for 12 hours in LB with carbenicillin while also estimating the number of transformed colonies by plating a serial dilution of transformation mixture as previously described^47^. We midiprepped the expanded transformations with ZymoPURE II Plasmid Midiprep (D4200).

### Building the Barcode-Promoter Dictionary

We introduced a unique 16-bp “plasmid barcode” adjacent to the enhancer sequence to allow us to determine from which promoter each transcript originated, which, together with the self-transcribed enhancer, allow us to map each transcript to a promoter-enhancer pair.

To build the map from 16-bp plasmid barcodes to promoters we PCR-amplified a fragment containing both the promoter and plasmid barcode from the plasmid library (98°C for 1 minute and 16 cycles of 98°C for 10 seconds, 66°C for 15 seconds, and 72°C for 25 seconds, ExP_P1_fwd_I2: AATGATACGGCGACCACCGAGATCTACAC[index-2]GGGAGGTATTGGACAGGC, ExP_P3_rev: CAAGCAGAAGACGGCATACGAGATGCATGGGTAGAAGTTAGCTGAAC) and sequenced the promoter position with paired-end reads (using custom sequencing primers ExP_P1_fwd_seq_R1: GAGTGAGCTCTCGTTCATCATGTGGGACATCAAGC, ExP_P2_rev_seq_R2: TGGTAGCAAGCTTAGATAAGGTGGACTCACTATGC) and the plasmid barcode with an index read (using custom sequencing primer ExP_fwd_BC_seq: GTCCCAATTCTTGTTGAATTAGATTGATCTAGAGCATGCA). We mapped these sequences to a specially constructed index of the promoter sequences using bowtie2 (X: -q --met-stderr -- maxins 2000 -p 4 --no-mixed --dovetail --fast). We dropped any BC-promoter pairs with singleton reads, then removed ambiguous pairings (more than one promoter for the same BC), and finally thresholded pairs with at least 5 reads to build the Barcode-Promoter dictionary.

### Cell Culture

We maintained cells at a density between 100,000 and 1,000,000 cells per ml in RPMI-1640 (Thermo Fisher Scientific) with 10% heat-inactivated FBS, 2 mM L-glutamine and 100 units per ml streptomycin and 100 mg ml−1 penicillin by diluting cells 1:8 in fresh medium every 3 days. Cell lines were regularly tested for mycoplasma.

### Library Transfection

We nucleofected 10 million K562 cells with 15µg of the ExP plasmid library in 100µL cuvettes with the Lonza 4D-Nucleofector using settings and protocols specified by the manufacturer for K562 cells (T-016). We pooled 5 nucleofections together during recovery to form 50 million cell biological transfection replicates and generated 4 replicates for a total of 200 million total cells. After 24 hours, we harvested the cells in Qiagen buffer RLT (79216) and proceeded with STARR-seq library preparation.

### STARR-seq Library Preparation

We proceeded with STARR-seq library preparation using an adapted protocol from Arnold 2013^4^. We split the 50 million-cell transfection replicates in half and extracted total RNA using 3 Qiagen RNeasy mini columns (74134), performing the on-column DNase step. We isolated polyA+ mRNA using the Qiagen Oligotex mRNA kit for the 1000 x 1000 ExP dataset (note this kit has been discontinued, we now use the Poly(A)Purist MAG kit from Thermo Fisher Scientific, AM1922). Following mRNA elution, we treated with TURBO DNase (Thermo Fisher Scientific, AM2238) in 100uL reactions at 37°C for 30 minutes, then added an additional 2uL of TURBO DNase and incubated at 37°C for 15 minutes. We purified the RNA following DNA digestion with Zymo RNA Clean & Concentrator 5 (R1013). We reverse transcribed the polyA+ mRNA using Thermo SuperScriptIV using the STARR_RT primer (CAAACTCATCAATGTATCTTATCATG) in 20uL reactions according to manufacturer’s recommendations. We included 1uL of ribonuclease inhibitor RNaseOUT (Invitrogen, 10777019). Following reverse transcription, we added 1uL of RNaseH (Thermo Fisher Scientific, EN0201) and incubated at 37°C for 20 minutes. We purified the cDNA with 1.8X volume of AMPure XP beads. We next selectively amplified the reporter transcript using intron-spanning junction primers with Q5 polymerase in 50uL reactions (98°C for 45 seconds and 15 cycles of 98°C for 15 seconds, 65°C for 30 seconds, and 72°C for 70 seconds, jPCR_fwd: TCGTGAGGCACTGGGCAG*G*T*G*T*C, jPCR_rev: CTTATCATGTCTGCTCGA*A*G*C, * = phosphorothioate bonds). Following purifications with 0.8X volume of AMPure XP beads, we performed a test final sequencing-ready PCR with a dilution of the junction PCR product to determine the optimal cycle number, then proceeded with the final PCR using Q5 polymerase in 50uL reactions (98°C for 45 seconds and ∼9 cycles of 98°C for 10 seconds, 65°C for 30 seconds, and 72°C for 30 seconds, ExP_GFP_fwd_I2: AATGATACGGCGACCACCGAGATCTACAC[index-2]GGCTTAAGCATGGCTAGCAAAG, ExP_P4_rev: CAAGCAGAAGACGGCATACGAGATTCATTCCATGGCATCTCACG. We purified the final libraries with 2 rounds of 0.8X volume of SPRISelect (Beckman Coulter, B23318).

### Alignment and counting of STARR-seq data

To characterize activity in the STARR-seq assay, we define “STARR-seq expression” for a given plasmid (corresponding to a particular promoter, enhancer, and plasmid barcode) as the expression of the reporter RNA transcript normalized to the abundance of that plasmid in the input DNA pool.

To quantify STARR-seq expression, we sequenced the library of RNA transcripts produced from replicate transfections (described above) along with the DNA input with paired-end reads (using custom sequencing primers ExP_P3_fwd_seq_R1: GAGTACTGGTATGTTCAGCTAACTTCTACCCATGC, ExP_P4_rev_seq_R2: TCATTCCATGGCATCTCACGACCTACTTAACTTGC) and the plasmid barcode with an index read (using custom sequencing primer ExP_fwd_BC_seq: GTCCCAATTCTTGTTGAATTAGATTGATCTAGAGCATGCA). We aligned reads for both the RNA and DNA libraries to the designed enhancer sequences using bowtie2 (bowtie2 options: -q --met 30 --met-stderr --maxins 2000 -p 16 --no-discordant --no-mixed --fast).

We counted reads separately from PCR replicates derived from each biological replicate of 50M transfected cells, and scaled each of the PCR replicates within a biological replicate such that they had the same total normalized counts, equal to the maximum across all PCR replicates. We combined counts into per-biological replicate counts for further processing. We used the BC-promoter dictionary to identify the promoter associated with each transcript. We used the same mapping and BC-promoter assignment process for DNA.

For subsequent analysis, we discarded plasmids that had fewer than 25 DNA reads or fewer than 1 RNA transcript reads from further processing.

### Computing technical reproducibility and influence of plasmid barcode sequences

To assess the technical reproducibility of ExP-STARR-seq, we first compared STARR-seq expression between biological replicate experiments. Specifically, we first combined data from biological replicates 1 & 2 and 3 & 4. Next, we correlated log_2_(RNA/DNA) for these groups before (Fig. 1b) and after (Fig. S1e) averaging across plasmid barcodes corresponding to the same enhancer-promoter pair.

We next assessed the variation between plasmids with the same enhancer and promoter sequences but different random 16-bp plasmid barcodes, because these 16 nucleotides of random sequence might contain transcription factor motifs or other sequences that affect STARR-seq expression. To do so, we combined data from all biological replicate experiments and created two “virtual replicates” for each enhancer-promoter pair by splitting the corresponding plasmid barcodes into two groups. For example, an enhancer-promoter pair with 6 plasmid barcodes was split into 2 virtual replicates each with 3 barcodes). We averaged log_2_ STARR-seq expression within enhancer-promoter pairs (across different barcodes) and correlated these virtual replicates. We compared versions of this analysis for increasing thresholds on the minimum number of barcodes in each virtual replicate (Fig. S1c,d).

### Estimating enhancer and promoter activities — naïve averaging approach

We sought to compare the intrinsic activities of different enhancer and promoter sequences in ExP STARR-seq — that is, the contribution of a given enhancer or promoter sequence to STARR-seq expression, relative to other sequences. We estimated enhancer activity and promoter activity in two ways: by a simple averaging method, and by fitting a multiplicative Poisson count model (see next section).

As a first approach to estimate promoter activity, we calculated, for each promoter sequence, the average log_2_ STARR-seq expression when that promoter is paired with random genomic sequences in the enhancer position (Fig. 1c). This quantity represents the “basal” or “autonomous” expression level of the promoter, in the absence of a strong activating sequence in the enhancer position.

As a first approach to estimate enhancer activity, we calculated, for each enhancer sequence, the average log_2_ STARR-seq expression of all pairs including that enhancer sequence (Fig. 1d).

As noted above, we fit this model on the set of plasmids with at least 25 DNA reads, and at least 1 RNA read. In addition, to reduce noise in our promoter and enhancer activity estimates, we required at least two separate plasmid barcodes per promoter-enhancer pair. These filters resulted in 604,268 promoter-enhancer pairs across 4,512,907 total unique plasmids (∼ 7.5 plasmids per pair) that were used to estimate promoter and enhancer activity.

In practice, this averaging method of calculating enhancer and promoter activity was inaccurate and biased, for several reasons. First, the averaging method does not consider the variance introduced by sampling & counting noise in sequencing, which is significant because many promoter-enhancer pairs have low RNA read counts. Second, the averaging method does not account for differences introduced due to missing data. In the 1000 enhancer x 1000 promoter data matrix, many entries are missing either due to low RNA counts (resulting from counting and sampling noise, or low expression) or due to low DNA counts (resulting from variation introduced in cloning the plasmid library). As a result of these factors, the averaging method produces biased (inflated) estimates of activity for weaker enhancer and promoter sequences because the expression of plasmids containing these sequences is more likely to drop below the threshold of detection given our sequencing depth (Fig. S2c-d).

Because this model explained the data well, we used this same model to estimate intrinsic enhancer and promoter activity.

### Estimating intrinsic enhancer and promoter activities — multiplicative model

We fit a count-based Poisson model to address the limitations of using a simple averaging approach to estimate intrinsic enhancer and promoter activities (see previous section), and to quantify the extent to which the ExP STARR-seq data can be explained by a simple multiplicative function of intrinsic enhancer and promoter activities. In this multiplicative model, all enhancers are assumed to activate all promoters by the same fold-change, without enhancer-promoter interaction terms.

Specifically, we estimate enhancer and promoter activities from ExP STARR-seq data by fitting the observed RNA read counts to a multiplicative function of observed DNA input read counts, intrinsic enhancer activity, and intrinsic promoter activity:

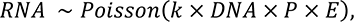

In this formula, *P* is the intrinsic promoter activity of promoter sequence p, *E* is intrinsic enhancer activity of enhancer sequence e, and *k* is a global scaling/intercept term that accounts for factors that control the relative counts of DNA and RNA such as sequencing depth.

We fit these parameters using block coordinate descent on the negative log-likelihood of the distribution above, initially fixing *k*=0, then alternatively optimizing (i) promoter activities while holding enhancer activities constant, and (ii) enhancer activities while holding promoter activities constant.

We then re-normalized enhancer activities and promoter activities by the mean activity of random genomic sequences, and adjusted the scaling factor *k* accordingly.

In practice, this model produces similar estimates to simply taking the mean value of an enhancer sequence across all promoters, and vice versa, but accounts for missing data points in the 1000×1000 matrix, and provides a more robust estimate for very weak enhancers or promoters, which produce relatively little RNA and are therefore difficult to measure in this STARR-seq experiment except when paired with a strong element in the other group (Fig. S2c-d).

### Computing and clustering residuals from the multiplicative model

We explored whether enhancer-promoter compatibility could explain variation in STARR-seq expression beyond that explained by the multiplicative model. To do so, we looked for shared behaviors between groups of promoters and enhancers by clustering them according to their residual error from the Poisson model described above.

For each enhancer-promoter pair, we used the Poisson model above to compute predicted RNA given the input DNA counts and estimates of intrinsic enhancer and promoter activities. We then compute a transformed residual as

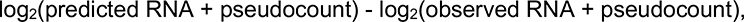

where pseudocount = 10 to stabilize variance of the estimates across the range of values for RNA^48^. We filtered to all enhancer-promoter pairs with at least two barcodes, and calculated the mean of the residuals across barcodes to form a (sparse) 1000×1000 matrix of residuals indexed by promoter and enhancers.

We clustered this matrix independently along rows and columns (treating missing pairs as having a residual of 0) using K-means with 3 clusters, labeling the clusters as 0,1, and 2 such that they had increasing mean activity estimates in the Poisson model. One cluster each of enhancers and promoters (E0 and P0) contained sequences that were missing many data points due to their weaker activity leading to dropout due to low RNA expression. The sparsity of data for the E0 and P0 clusters prevented accurate characterization of compatibility, and so we excluded these clusters from subsequent analysis.

### Assessing reproducibility of the clusters

We evaluated whether the clustering we observed in the residuals was a general trend of the data, or an artifact of a few promoters or enhancers. To test this possibility, we randomly downsampled the residual matrix to 25% of promoters and 25% of enhancers (6.25% of the total data) 100 times, and clustered the subsets. We found that the original (full-data) cluster identity of a promoter or enhancer predicted the downsampled cluster with greater than 80% accuracy (Fig. S3g).

### Estimating enhancer activity with specific promoter classes, and promoter responsiveness to specific enhancer classes

We evaluated whether certain promoters were more responsive when paired with different enhancer classes, and whether certain enhancers had more activity when paired with promoters from different classes (Fig. 3c,d).

To explore differences in enhancer activity when paired with different promoter classes, we fit the Poisson model (described above) separately to two different subsets of the data: (i) all enhancer sequences paired with P1 or genomic background promoter sequences (yielding an estimate of the activity of an enhancer sequence on a P1 promoter), and (ii) all enhancer sequences paired with P2 or genomic background promoter sequences (yielding an estimate of the activity of an enhancer sequence on a P2 promoter).

Similarly, to estimate promoter responsiveness to either E1 or E2 enhancers, we fit the Poisson model to the subsets: (iii) all promoters paired with E1 or genomic background enhancer sequences (yielding an estimate of the responsiveness of a promoter sequence to E1 enhancers), and (iv) all promoters paired with E2 or genomic background enhancer sequences (yielding an estimate of the responsiveness of a promoter sequence to E2 enhancers).

We used the genomic background promoter sequences to set a common baseline.

### Annotating enhancer and promoter sequences with genomic features and sequence motifs

To annotate enhancer and promoter sequences with features of transcription factor (TF) binding of the corresponding genomic elements, we downloaded list of Human TF ChIP-seq narrowpeak files from the ENCODE Project^21^, and annotated each enhancer or promoter sequence with the maximum signalValue column for any overlapping peak (or 0 signal, for no overlap). We then compared the fold-change in signal between classes of sequences (Fig. 4d, Fig. S5a, Table S9).

To annotate enhancer and promoter sequences with transcription factor motifs, we used FIMO^49^ (default parameters, including *p*-value threshold of 10^-6^) to identify matches for HOCOMOCO v11 CORE motifs^50^. We then compared the fold-change in motif counts between classes of sequences (Fig. S6f, Table S5, Table S7).

For comparing features between E1 and E2 enhancers, we compared motif, ChIP-seq, and other features between the E1 and E2 enhancer sequences that overlapped the summit of a DNase peak.

For analyzing the proportion of P2 promoters bound by various factors, we defined “strongly bound” as having ChIP-seq signal greater than 20% of maximum ChIP-seq signal among P1 and P2 promoters.

### Comparison of CRISPR-derived regulatory elements for P1 vs P2 promoters

To compare the number and effect sizes of genomic regulatory elements for P1 and P2 promoters, we analyzed CRISPRi tiling screens from previous studies that perturbed all DNase accessible sites around selected genes^22, 26, 27^. We counted the number of activating distal regulatory elements — *i.e.*, distal, non-promoter DNase accessible sites whose perturbation led to a significant reduction in gene expression (Fig. 4c). We also compared the effect sizes on gene expression for these same activating distal regulatory elements (Fig. S5c, Fig. S6b).

### Luciferase assays

We tested the ability of each of 7 large *MYC* enhancer fragments to activate the promoters of 3 genes in the *MYC* locus — *MYC*, *PVT1*, and *CCDC26* — using a classic plasmid luciferase-based enhancer assay. The 7 *MYC* enhancers were defined as the 1.0-2.2 kb sequences identified in our previous *MYC* proliferation-based CRISPRi screen^27^, and a 1 kb bacterial plasmid sequence was used as a negative control sequence. We cloned promoter fragments into plasmids in combination with each of these sequences. The promoter fragments corresponded to the dominant transcription start site of each gene in K562 cells (as determined by CAGE). For each of *PVT1* and *CCDC26* — which do not appear to be regulated by most of the 7 *MYC* enhancers in the genome — we cloned two promoter fragments of different lengths to determine if nearby sequences might encode biochemical specificity. We designed an insertion site ∼1 kb upstream of the promoter in the plasmid for inserting each enhancer sequence (Fig. S2g), and we flanked this region with polyadenylation signals in either direction to avoid measuring luciferase activity driven from transcripts initiating from the enhancer elements themselves. Luciferase assays using the Dual-Luciferase Reporter Assay (Promega) were performed as previously described^27^ in biological triplicate. For each experiment, we calculated the fold-change in luciferase signal (Firefly / Renilla) for enhancer versus negative control (Fig. S2i).

### Assessing the cell-type specificity of E1 and E2 enhancers

We tested whether E1 and E2 enhancer sequences from ExP STARR-seq overlapped elements predicted to act as enhancers by the ABC model in K562 cells or in 128 other cell types and tissues. To do so, we intersected the E1 and E2 enhancer sequences with the ∼200-bp regions predicted by the ABC model to act as enhancers for at least 1 nearby expressed gene, as previously defined^51^. The ABC enhancer-gene predictions from this previous study^51^ are available at https://www.engreitzlab.org/resources/.

### Aligning promoters by transcription start site

For each 264-bp promoter sequence, we defined the primary transcription start site (TSS) as the nucleotide with the highest stranded 5’ signal in GRO-Cap data in K562 cells (GSM1480321)^52^. This primary TSS position was used for plotting genomic signals relative to TSS and in analyses of motif positioning (*e.g.*, for GABPA and YY1).

### Analysis of motif position relative to TSS

We used FIMO^49^ to scan for HOCOMOCO motifs in promoters including for GABPA (GABPA_HUMAN.H11MO.0.A), YY1 (TYY1_HUMAN.H11MO.0.A), and the TATA box (TBP_HUMAN.H11MO.0.A). We reported positional preferences as the distance between the primary transcription start site from GRO-cap (see above) and the center of the motif. For example, GABPA, the most common position was –10 relative to the TSS (*i.e.* with the second ‘G’ in the core ‘GGAA’ motif located at position –10).

### Hybrid selection STARR-seq (HS-STARR-seq) to measure enhancer activity for millions of genomic sequences

We conducted two STARR-seq experiments to measure the enhancer activity of millions of long genomic sequences tiling across human enhancer and promoter sequences. To generate these tiling sequences, we used a hybrid selection strategy, similar to previous approaches^53^. Specifically, we purified genomic DNA from K562 cells, tagmented DNA using Tn5 and gel size selection to a size range of approximately 300-700 bp (Fig. S8c), and conducted hybrid selection using RNA probes as previously described^54^ targeting either (i) all gene promoters (“HS promoter pool”) or (ii) all accessible elements (“HS accessible element pool”) in K562 cells (see Table S10 and Table S11 for probe sequences). We amplified these sequences using primers including a UMI (CapStarrFa_N10 primer: tagatTGAtCTAGAGCATGCACCGGCAAGCAGAAGACGGCATACGAGATNNNNNNNNNNATGTCTCGTGGGCTCGGAGATGT and CapStarrR primer: CGAAGCGGCCGGCCGAATTCGTCGATCGTCGGCAGCGTCAGATGTG) and cloned these selected sequences into the hSTARR-seq_ori vector^8^, which uses the bacterial origin of replication (ORI) sequence as the promoter for the reporter transcript, by Gibson assembly. In the final HS promoter and accessible element Pools, 9% and 12% of fragments mapped to their intended targets, respectively, and each element was tiled by a median of 20 and 55 sequences. We conducted the rest of the STARR-seq experiment as described above, transfecting 50 million cells per replicate for each of 4 replicates.

We sequenced the input DNA libraries to a depth of 880 million and 810 million reads (promoter and accessible element pools, respectively), and the RNA libraries to a depth of 1.1 billion reads (both pools). We aligned reads to the hg19 genome using bowtie2 (options: -q --met-stderr -- maxins 1000 -p 4 --no-discordant --no-mixed). We discarded fragments with fewer than 25 aligned DNA reads. Biological replicates were highly correlated (*R*^2^ = 0.92 and 0.91 for promoter and accessible element pools) (Fig. S8b).

We analyzed this data by computing a log_2_ activity per fragment equal to the log_2_(RNA / DNA). and correcting for a fragment-length bias. We noted that STARR-seq expression was highly inversely correlated with the length of the enhancer sequence, even among random genomic fragments that did not overlap putative regulatory elements, which could result from biases in library preparation and sequencing. To adjust for this, we fit a linear regression (separately for the two pools) and subtracted this regression from the log_2_(RNA / DNA) activity to give a bias-corrected activity. We then correlated motifs with bias-corrected activity. To estimate enhancer activity of promoters from the ExP, we found HS-STARR-seq fragments that overlapped at least 90% of an ExP promoter and averaged their activity scores.

## References

1. Banerji, J., Rusconi, S. & Schaffner, W. Expression of a beta-globin gene is enhanced by remote SV40 DNA sequences. Cell 27, 299–308 (1981).

2. Banerji, J., Olson, L. & Schaffner, W. A lymphocyte-specific cellular enhancer is located downstream of the joining region in immunoglobulin heavy chain genes. Cell 33, 729–740 (1983).

3. Melnikov, A. et al. Systematic dissection and optimization of inducible enhancers in human cells using a massively parallel reporter assay. Nat. Biotechnol. 30, 271–277 (2012).

4. Arnold, C. D. et al. Genome-wide quantitative enhancer activity maps identified by STARR-seq. Science 339, 1074–1077 (2013).

5. Kermekchiev, M., Pettersson, M., Matthias, P. & Schaffner, W. Every enhancer works with every promoter for all the combinations tested: could new regulatory pathways evolve by enhancer shuffling? Gene Expr. 1, 71–81 (1991).

6. Tewhey, R. et al. Direct Identification of Hundreds of Expression-Modulating Variants using a Multiplexed Reporter Assay. Cell 172, 1132–1134 (2018).

7. Klein, J. C. et al. A systematic evaluation of the design and context dependencies of massively parallel reporter assays. Nat. Methods 17, 1083–1091 (2020).

8. Muerdter, F. et al. Resolving systematic errors in widely used enhancer activity assays in human cells. Nat. Methods 15, 141–149 (2018).

9. Nguyen, T. A. et al. High-throughput functional comparison of promoter and enhancer activities. Genome Res. 26, 1023–1033 (2016).

10. Emami, K. H., Navarre, W. W. & Smale, S. T. Core promoter specificities of the Sp1 and VP16 transcriptional activation domains. Mol. Cell. Biol. 15, 5906–5916 (1995).

11. Ohtsuki, S., Levine, M. & Cai, H. N. Different core promoters possess distinct regulatory activities in the Drosophila embryo. Genes Dev. 12, 547–556 (1998).

12. Emami, K. H., Jain, A. & Smale, S. T. Mechanism of synergy between TATA and initiator: synergistic binding of TFIID following a putative TFIIA-induced isomerization. Genes Dev. 11, 3007–3019 (1997).

13. Butler, J. E. F. Enhancer-promoter specificity mediated by DPE or TATA core promoter motifs. Genes & Development vol. 15 2515–2519 (2001).

14. Yean, D. & Gralla, J. Transcription reinitiation rate: a special role for the TATA box. Molecular and Cellular Biology vol. 17 3809–3816 (1997).

15. Wefald, F. C., Devlin, B. H. & Williams, R. S. Functional heterogeneity of mammalian TATA-box sequences revealed by interaction with a cell-specific enhancer. Nature 344, 260–262 (1990).

16. Zabidi, M. A., Arnold, C. D., Schernhuber, K. & Pagani, M. Enhancer–core-promoter specificity separates developmental and housekeeping gene regulation. Nature (2015).

17. Arnold, C. D. et al. Genome-wide assessment of sequence-intrinsic enhancer responsiveness at single-base-pair resolution. Nat. Biotechnol. 35, 136–144 (2017).

18. Haberle, V. et al. Transcriptional cofactors display specificity for distinct types of core promoters. Nature 570, 122–126 (2019).

19. van Arensbergen, J., van Steensel, B. & Bussemaker, H. J. In search of the determinants of enhancer–promoter interaction specificity. Trends in Cell Biology vol. 24 695–702 (2014).

20. Li, X. & Noll, M. Compatibility between enhancers and promoters determines the transcriptional specificity of gooseberry and gooseberry neuro in the Drosophila embryo. EMBO J. 13, 400–406 (1994).

21. ENCODE Project Consortium. An integrated encyclopedia of DNA elements in the human genome. Nature 489, 57–74 (2012).

22. Fulco, C. P. et al. Activity-by-contact model of enhancer-promoter regulation from thousands of CRISPR perturbations. Nat. Genet. 51, 1664–1669 (2019).

23. Wall, L., deBoer, E. & Grosveld, F. The human beta-globin gene 3’ enhancer contains multiple binding sites for an erythroid-specific protein. Genes Dev. 2, 1089–1100 (1988).

24. Tuan, D. Y., Solomon, W. B., London, I. M. & Lee, D. P. An erythroid-specific, developmental-stage-independent enhancer far upstream of the human ‘beta-like globin’ genes. Proc. Natl. Acad. Sci. U. S. A. 86, 2554–2558 (1989).

25. Thakore, P. I. et al. Highly specific epigenome editing by CRISPR-Cas9 repressors for silencing of distal regulatory elements. Nat. Methods 12, 1143–1149 (2015).

26. Klann, T. S. et al. CRISPR-Cas9 epigenome editing enables high-throughput screening for functional regulatory elements in the human genome. Nat. Biotechnol. 35, 561–568 (2017).

27. Fulco, C. P. et al. Systematic mapping of functional enhancer-promoter connections with CRISPR interference. Science 354, 769–773 (2016).

28. Liu, Y. et al. Functional assessment of human enhancer activities using whole-genome STARR-sequencing. Genome Biol. 18, 219 (2017).

29. Haberle, V. & Stark, A. Eukaryotic core promoters and the functional basis of transcription initiation. Nat. Rev. Mol. Cell Biol. 19, 621–637 (2018).

30. Lenhard, B., Sandelin, A. & Carninci, P. Metazoan promoters: emerging characteristics and insights into transcriptional regulation. Nat. Rev. Genet. 13, 233–245 (2012).

31. Fan, K., Moore, J. E., Zhang, X.-O. & Weng, Z. Genetic and epigenetic features of promoters with ubiquitous chromatin accessibility support ubiquitous transcription of cell-essential genes. Nucleic Acids Res. 49, 5705–5725 (2021).

32. Xi, H. et al. Identification and characterization of cell type-specific and ubiquitous chromatin regulatory structures in the human genome. PLoS Genet. 3, e136 (2007).

33. Landolin, J. M. et al. Sequence features that drive human promoter function and tissue specificity. Genome Res. 20, 890–898 (2010).

34. Weingarten-Gabbay, S. et al. Systematic interrogation of human promoters. Genome Res. 29, 171–183 (2019).

35. Sahu, B., Hartonen, T., Pihlajamaa, P., Wei, B. & Dave, K. Sequence determinants of human gene regulatory elements. bioRxiv (2021).

36. Yu, M. et al. GA-binding protein-dependent transcription initiator elements. Effect of helical spacing between polyomavirus enhancer a factor 3(PEA3)/Ets-binding sites on initiator activity. J. Biol. Chem. 272, 29060–29067 (1997).

37. Curina, A. et al. High constitutive activity of a broad panel of housekeeping and tissue-specific cis-regulatory elements depends on a subset of ETS proteins. Genes Dev. 31, 399–412 (2017).

38. van Arensbergen, J. et al. Genome-wide mapping of autonomous promoter activity in human cells. Nat. Biotechnol. 35, 145–153 (2017).

39. Hong, C. K. Y. & Cohen, B. A. Genomic environments scale the activities of diverse core promoters. doi:10.1101/2021.03.08.434469.

40. Maricque, B. B., Chaudhari, H. G. & Cohen, B. A. A massively parallel reporter assay dissects the influence of chromatin structure on cis-regulatory activity. Nat. Biotechnol. (2018) doi:10.1038/nbt.4285.

41. Chiang, C. M. & Roeder, R. G. Cloning of an intrinsic human TFIID subunit that interacts with multiple transcriptional activators. Science 267, 531–536 (1995).

42. Austen, M., Lüscher, B. & Lüscher-Firzlaff, J. M. Characterization of the transcriptional regulator YY1. The bipartite transactivation domain is independent of interaction with the TATA box-binding protein, transcription factor IIB, TAFII55, or cAMP-responsive element-binding protein (CPB)-binding protein. J. Biol. Chem. 272, 1709–1717 (1997).

43. Sucharov, C., Basu, A., Carter, R. S. & Avadhani, N. G. A novel transcriptional initiator activity of the GABP factor binding ets sequence repeat from the murine cytochrome c oxidase Vb gene. Gene Expr. 5, 93–111 (1995).

44. Carter, R. S. & Avadhani, N. G. Cooperative binding of GA-binding protein transcription factors to duplicated transcription initiation region repeats of the cytochrome c oxidase subunit IV gene. J. Biol. Chem. 269, 4381–4387 (1994).

45. Usheva, A. & Shenk, T. YY1 transcriptional initiator: protein interactions and association with a DNA site containing unpaired strands. Proc. Natl. Acad. Sci. U. S. A. 93, 13571–13576 (1996).

46. Larsson, A. J. M. et al. Genomic encoding of transcriptional burst kinetics. Nature 565, 251–254 (2019).

47. Wang, T., Lander, E. S. & Sabatini, D. M. Large-Scale Single Guide RNA Library Construction and Use for CRISPR-Cas9-Based Genetic Screens. Cold Spring Harb. Protoc. 2016, db.top086892 (2016).

48. Anscombe, F. J. THE TRANSFORMATION OF POISSON, BINOMIAL AND NEGATIVE-BINOMIAL DATA. Biometrika vol. 35 246–254 (1948).

49. Grant, C. E., Bailey, T. L. & Noble, W. S. FIMO: scanning for occurrences of a given motif. Bioinformatics 27, 1017–1018 (2011).

50. Kulakovskiy, I. V. et al. HOCOMOCO: towards a complete collection of transcription factor binding models for human and mouse via large-scale ChIP-Seq analysis. Nucleic Acids Res. 46, D252–D259 (2018).

51. Nasser, J. et al. Genome-wide enhancer maps link risk variants to disease genes. Nature 593, 238–243 (2021).

52. Core, L. J. et al. Analysis of nascent RNA identifies a unified architecture of initiation regions at mammalian promoters and enhancers. Nat. Genet. 46, 1311–1320 (2014).

53. Vanhille, L. et al. High-throughput and quantitative assessment of enhancer activity in mammals by CapStarr-seq. Nat. Commun. 6, 6905 (2015).

54. Engreitz, J. M. et al. Local regulation of gene expression by lncRNA promoters, transcription and splicing. Nature 539, 452–455 (2016).

